# Collecting, detecting and handling non-wear intervals in longitudinal light exposure data

**DOI:** 10.1101/2024.12.23.627604

**Authors:** Carolina Guidolin, Johannes Zauner, Steffen Lutz Hartmeyer, Manuel Spitschan

## Abstract

In field studies using wearable light loggers, participants often need to remove the devices, resulting in non-wear intervals of varying and unknown duration. Accurate detection of these intervals is an essential step during data pre-processing. Here, we deployed a multi-modal approach to collect non-wear time during a longitudinal light exposure collection campaign and systematically compare non-wear detection strategies. Healthy participants (n=26; mean age 28±5 years, 14F) wore a near-corneal plane light logger for one week and reported non-wear events in three ways: pressing an “event marker” button on the light logger, placing it in a black bag, and using an app-based Wear log. Wear log entries, checked twice daily, served as ground truth for non-wear detection, showing that non-wear time constituted 5.4±3.8% (mean±SD) of total participation time. Button presses at the start and end of non-wear intervals were identified in >85.4% of cases when considering time windows beyond one minute for detection. To detect non-wear intervals based on black bag use and lack of motion, we employed an algorithm detecting clusters of low illuminance and clusters of low activity. Performance was higher for illuminance (F1=0.78) than activity (F1=0.52). Light exposure metrics derived from the full dataset, a dataset filtered for non-wear based on self-reports, and a dataset filtered for non-wear using the low illuminance clusters detection algorithm showed minimal differences. Our results highlight that while non-wear detection may be less critical in high-compliance cohorts, systematically collecting and detecting non-wear intervals is feasible and important for ensuring robust data pre-processing.

## Introduction

### The use of light loggers to measure personal light exposure

Ocular light exposure in everyday life is increasingly being recognised for its potential to influence human physical and mental wellbeing, including cognitive functions, sleep, circadian health, and eye health (Brown et al., 2022; Muralidharan et al., 2021). In this framework, measuring light exposure in the real world is essential to understand what light exposure individuals receive in their daily lives and the cultural, geographical, architectural, and behavioural contexts underlying their exposure patterns (Biller et al., 2024; Guidolin et al., 2024). The measurement of light exposure in free-living conditions is largely accessible in research contexts due to the availability of light loggers, that is, wearable devices recording light exposure over extended periods of time (Hartmeyer et al., 2022; van Duijnhoven et al., 2024). Depending on the intended measurement location, light loggers come in various form factors, including wrist-watches (wrist level measurements), pendants or brooches (chest level measurements), and removable attachments to non-prescription glasses frames (near-corneal plane level measurement) (Hartmeyer et al., 2022). The collected light exposure data can either be directly analysed to classify exposure patterns or also aggregated and summarised in light exposure metrics, which can in turn be associated with health-related outcomes of interest (Spitschan et al., 2022). The versatility of wearable light loggers has led to their use for different research applications (for a detailed review of studies using light loggers in the context of circadian rhythms, alertness, cognitive performance and mood, see Hartmeyer et al., 2022). Most commonly, light loggers have been used in observational field studies aiming to capture the effects of light exposure on sleep (Mead et al., 2023), alertness (Didikoglu et al., 2023), and metabolic function (Reid et al., 2014). Furthermore, light loggers integrated into actigraphy devices have been used in large-scale epidemiological studies to assess light exposure as a potential risk factor for disease prevalence observed at follow-up (Burns et al., 2023; Windred et al., 2024; Windred et al., 2024). Finally, a few studies have employed light loggers in clinical settings to monitor compliance with prescribed light regimes (Sit et al., 2018) or in occupational health to assess whether employees meet target light levels during working hours (Peeters et al., 2020).

### Data quality challenges in pre-processing light exposure data: non-wear intervals

Light logging also poses challenges to researchers interested in extracting light metrics from measured light levels, including data quality assessment. The raw wearable data is usually exported as a singular data file containing a time series of light levels and other logged variables, such as activity and external temperature, depending on the individual device (van Duijnhoven et al., 2024). These light data can contain artefacts and data recorded while the device was not worn (Hartmeyer et al., 2022). In fact, in field studies using light loggers, participants may need to remove the devices, such as when taking a shower, or during certain physical activities and sports. Moreover, participants may forget to wear the device or choose to remove it for miscellaneous reasons (Stefani et al., 2024). These periods of non-wear result in invalid data included in the collected datasets. To date, best practices for data cleaning pipelines have not been systematically investigated yet, and implemented strategies are seldom reported in the methods section of research papers (Hartmeyer et al., 2022). Hence, it is currently unknown whether non-wear intervals impact the derived light exposure metrics. It is important to note that the challenge of detecting non-wear time is not exclusive to wearable light loggers, but common across other research wearable devices, such as wrist-worn actimeters used for measuring rest and activity rhythms in field settings. In actimetry research, a variety of algorithms leveraging skin temperature, skin conductance, and (lack of) wearable movement have been developed to detect non-wear intervals (Choi et al., 2012; Pilz et al., 2022; Van Der Donckt et al., 2024). However, skin temperature and conductance might not be available in light loggers, or it might not be a useful measure to detect non-wear for light loggers that are not placed directly on the participants’ skin. Furthermore, contrary to movement in actimetry, non-wear time of light loggers is not necessarily characterised by a lack of light signal, since leaving the device on a window sill could lead to light levels similar to those of someone sitting in the same room and wearing the device. Some studies have addressed this issue by relying on intervals of device inactivity to identify invalid periods, using cut-off lengths ranging from 5 to 120 minutes of inactivity (Hartmeyer et al., 2022). Other approaches to detect non-wear intervals include relying on self-reported non-wear times, combining minimal variability in light signal and activity to detect non-wear time, and detecting stability in device tilt angles (Daugaard et al., 2017; Price et al., 2019; Stefani et al., 2024; Van der Maren et al., 2018). However, whether these methods were data-driven or involved manual inspection of participants’ files often remains unclear. Furthermore, even when data-driven techniques are employed, no study thus far has systematically analysed the performance of algorithmic approaches using a ground truth for comparison.

### The current study

The current study aims to describe and compare strategies for collecting, detecting and handling non-wear time of light exposure data using a high-quality ground truth for non-wear time intervals. To achieve this, during a week-long ambulatory light exposure study, we first collected non-wear information from three independent sources: participants’ self-reports, button presses on the light logger, and placement of the light logger in an opaque black cloth bag during non-wear time. Self-reported entries were checked twice daily to ensure high data quality and used as ground truth for non-wear interval detection. Second, we examine how non-wear detection varies based on non-wear sources. As part of this analysis, we employ an algorithm to detect clusters of low illuminance resulting from black bag use during non-wear, and clusters of low activity resulting from lack of motion during non-wear. We discuss its performance and how it could be leveraged to develop robust non-wear detection strategies further. Lastly, we assessed the impact of non-wear data on the accuracy of daily light exposure metrics.

## Methods and materials

### Procedure

This study aimed to identify individual and contextual factors contributing to light exposure patterns in free-living conditions, and a detailed protocol of the study was previously described (Guidolin et al., 2024). The analyses related to this study objective will be reported in a separate publication. In the current paper, we use the data collected during this data collection campaign to investigate collection, detection and handling of non-wear behaviour. Healthy participants aged 18 to 55 years were recruited to participate in a study on light exposure in everyday life lasting for a week (Monday to Monday) in Tübingen, Germany (48.5216° N, 9.0576° E) between August and November 2023. At time of recruitment, we aimed for a convenience sample of N=24 due to resource and time constraints. Later systematic analyses of sample size and statistical power requirements for light exposure studies in the field have validated that a sample size of N=24 provides a 80% statistical power for most light exposure metrics (Zauner et al., 2024).

### Near-corneal plane light logging

Participants wore a small commercial, multispectral 10-channel light logger (ActLumus, Condor Instruments, São Paolo, Brazil) able to measure photopic illuminance and melanopic equivalent daylight illuminance (mEDI). The device further contains an accelerometer, temperature sensor, and an ‘event’ button accessible to the participants. The light logger was inserted into a custom-made 3D-printed holder, which was in turn attached to the frame of non-prescription glasses (**Figure 1**). The glasses were available in sizes S, M and L, and provided by the brand Firmoo (**Figure 1** and **Supplementary Materials**). This set-up was chosen to ensure light exposure is measured at the near-corneal plane, which is less susceptible to clothing occlusion than wrist and chest measurements and a more accurate measurement point for measuring the physiological role of ocular light exposure. The sampling interval of the light logger was set to 10 seconds to achieve highly temporally resolved light exposure data. To ensure clarity in our explanations to participants and in this document, we referred to the combination of the non-prescription glasses and the mounted ActLumus light logger together as “light glasses”. Participants were instructed to wear the light glasses as much as possible throughout their participation in the study and only remove them in case of sleep, water sports, contact sports and during showering. Participants were compensated €10 for each day of wearing the light glasses for at least 80% of their waking hours.

**Figure 1.**
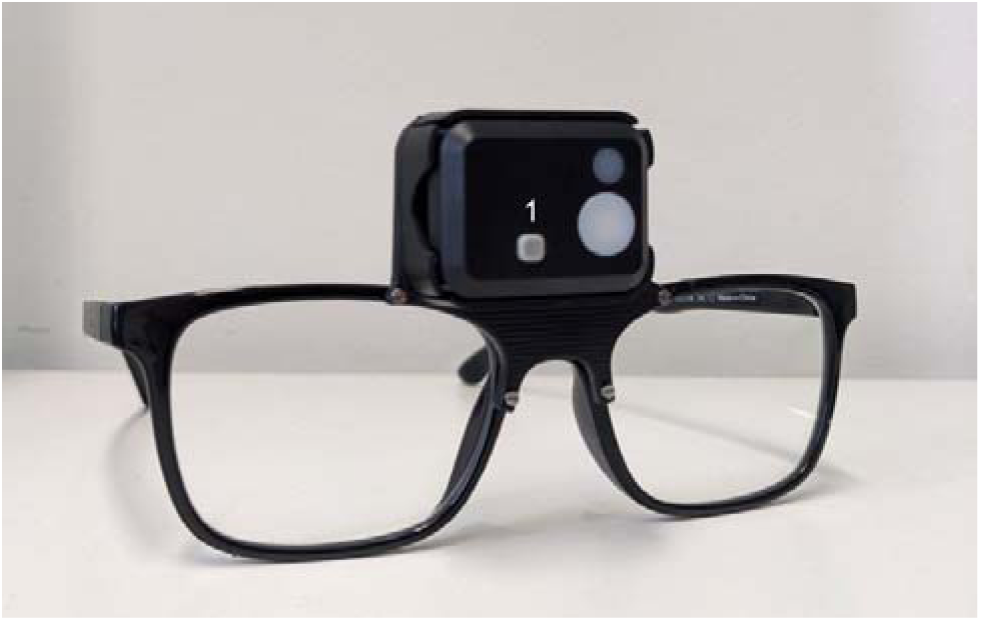
Light glasses worn by participants throughout the study. *Note.* The light glasses comprised non-prescription glasses and the ActLumus light logger mounted centrally on the bridge of the glasses frame. (1) Button (event marker) on the light logger.

### Logging and coding of non-wear events

Throughout the week, participants reported (non-)wear events, which included taking the light glasses off, placing the light glasses back on, and taking the light glasses off before sleep. These events were logged in three ways: (1) Pressing the event button on the light logger, (2) placing the light glasses in an opaque black-cloth bag (UVEX microfibre bag for spectacles) for the entire duration of non-wear, and (3) manually entering the time of this event in a Wear log on the mobile app MyCap (Harris et al., 2022). Participants were instructed as follows. When taking the light glasses off, they were to press the event button on the light glasses and then place them in the black bag provided. Participants were then to log this action in the app-based Wear log as “Taking the light glasses off”. Likewise, when putting the light glasses back on, they were instructed to take them out of the black bag, press the event button and log this action in the Wear log as “Putting the light glasses on”. In the Wear log, participants also had to indicate whether the light glasses were actually placed in the black bag. If they forgot the black bag, they had to describe where they placed the light glasses instead. When taking the light glasses off before sleep, participants were to place them facing upwards on a bedside table or flat surface near their bed and to log this action in the Wear log as “Taking the light glasses off before sleep and placing them on a nightstand”. Each Wear log entry included an automatic timestamp of the moment this action was performed. If they forgot to log an event, participants were allowed to retrospectively log any of the possible Wear log events as a “past event”. In this case, they were instructed to try their best to remember and retrospectively report the times of these actions.

This strategy for collecting non-wear intervals in three ways led to three independent non-wear sources: (1) A binary outcome for each button press (0/1) automatically added in the raw ActLumus data file, (2) clusters of near zero-lux illuminance levels while the device was placed in the black bag, and (3) a timestamp for each Wear log entry within the MyCap record (**Figure 2**). Among these three sources of non-wear information, the Wear log entries were deemed the cleanest based on continuous data quality assurance checks (see *Continuous non-wear data quality assurance* in *Data pre-processing*), and thus considered as the ground truth for our analyses (see *Analysis of button presses* and *Analysis of low illuminance clusters and low activity clusters*).

**Figure 2.**
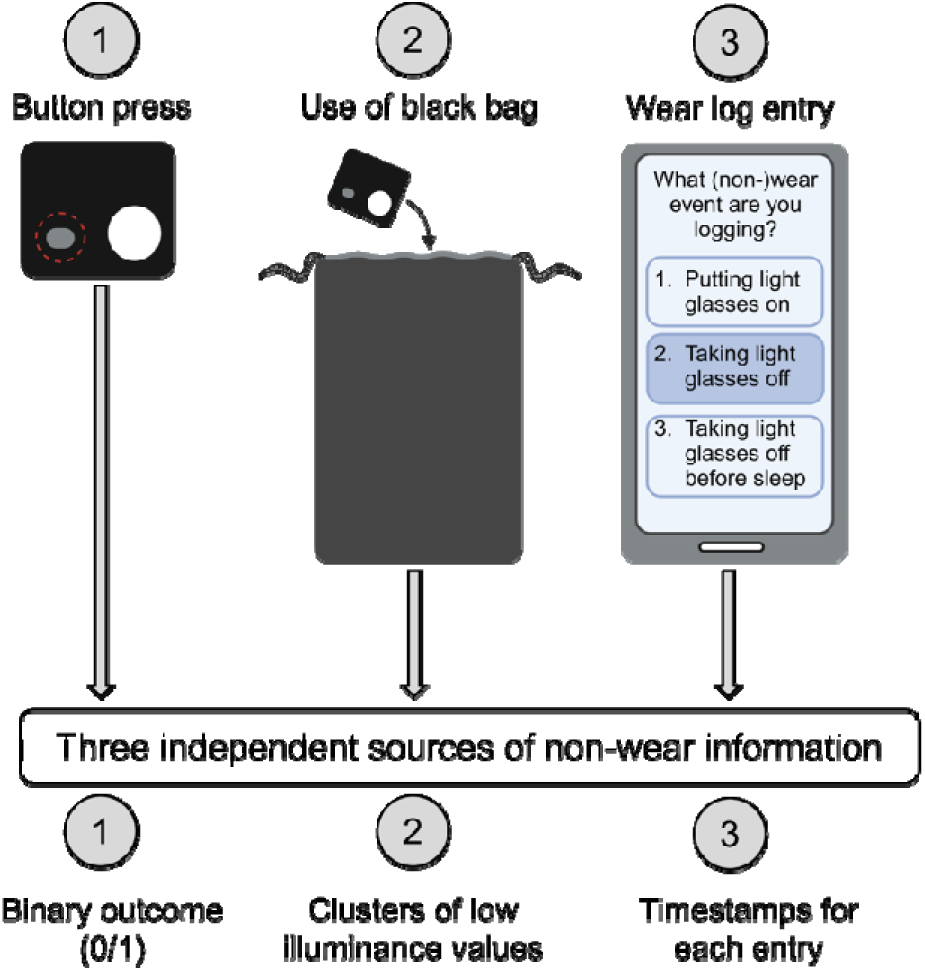
Schematic diagram of the non-wear intervals collection strategy. *Note.* Three independent methods of non-wear intervals were collected, leading to three independent source of non-wear information to base our analytic strategy on. This image was partially created using *BioRender.com*.

### Data analysis

All processing steps as well as all statistical analyses were performed in R statistical software (version 4.4.5) (R Core Team, 2025), using the packages “tidyverse” (version 2.0.0) (Wickham et al., 2019),“LightLogR” (version 0.9.2) (Zauner et al., 2025), “stats” (base package, version 4.5.0; R Core Team, 2025) and “rstatix” (version 0.7.2; Alboukadel, 2024).

#### Data pre-processing

Pre-processing of multi-modal data sets, such as the one used in the current study, is a time-consuming step. It demands a principled, systematic and reproducible approach. While scientific publications often report “data cleaning” and/or “data transformation” steps in the Methods section, the entire pre-processing pipeline is seldom reported in a detailed, systematic way (Hartmeyer et al., 2022) that allows other investigators to reproduce these analyses. This leads to multiple similar yet likely not equivalent ways of approaching the same type of data. In the following paragraphs, we summarise the pre-processing approach taken in the current study for handling each of the three non-wear sources to inspire other researchers working with wearable light loggers and facing similar challenges. Additional detailed information on our pre-processing strategy can be found in the **Supplementary Materials**.

Our pre-processing pipeline can be divided into two steps: (1) continuous non-wear data quality assurance, taking place during the study, and (2) data cleaning and transformation, taking place after the end of data collection.

##### Continuous non-wear data quality assurance

Throughout the study, we performed data quality assurance as participants were enrolled in the study. This step of our pre-processing pipeline was only applied to the Wear log entries, as information about button presses and the use of the black bag, as indicated by low illuminance values, was only available upon study completion. A researcher checked participants’ incoming Wear log entries at least twice daily, once in the morning and once in the late afternoon. This quality assurance was performed using the online platform REDCap (Research Electronic Data Capture; Harris et al., 2009), which was used for survey data collection and provides the backend with MyCap (Harris et al., 2022). The goal of this continuous check was to ensure that the collected data could be trusted and increase participants’ adherence to the protocol, thus ensuring high data quality throughout the study. For each participant, a spreadsheet was used to log the manual quality assurance and colour-coding valid Wear log entries as green and invalid (unclear and/or missing) entries as red. **Figure 3** illustrates how invalid entries were identified and adjusted, and further examples of invalid entries are provided in the **Supplementary Materials**.

**Figure 3.**
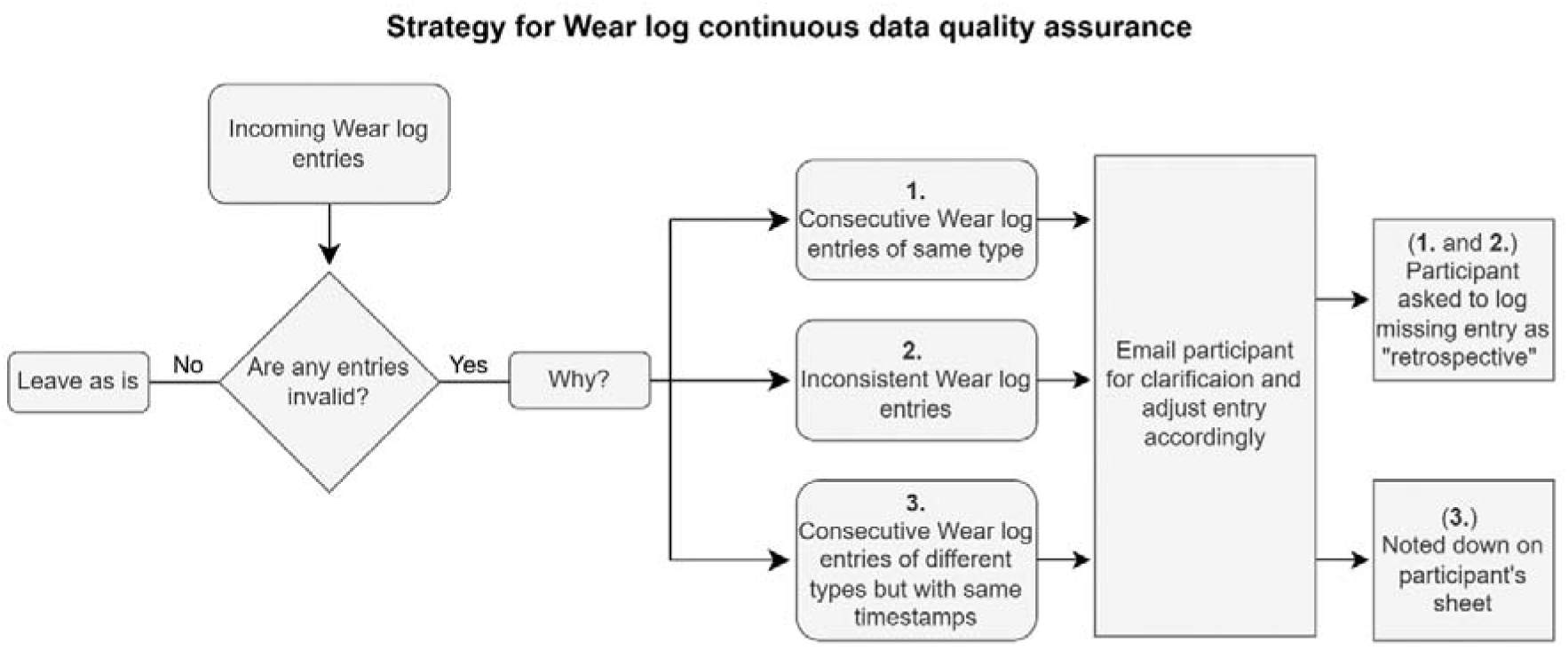
Flowchart for continuous quality assurance of Wear log entries performed throughout the study. *Note.* Rectangles indicate processes, rhombuses are used for yes/no branching, and rectangles with smoothed corners are used for descriptions.

##### Data cleaning of Wear log entries

A detailed, step-by-step description of the data cleaning and transformation approach undertaken for the Wear log entries is provided as a flowchart in the Supplementary Materials (**Figure S1**). The goal of the Wear log pre-processing pipeline was to identify intervals that participants spent wearing the light glasses (Wake □on□), not wearing the light glasses while awake (Wake □off□), and not wearing the light glasses during sleep (Sleep □off□). To further examine the validity of the Wear log entries beyond the manual quality assurance procedures performed during the study (see above under *Continuous non-wear data quality assurance*), two additional types of quality checks were performed:

1. Quality check #1: Identifying identical timestamps for two consecutive Wear log states, i.e. identical and consecutive Datetime values.
2. Quality check #2: Identifying consecutive identical Wear log states, i.e. identical and consecutive Wake □on□, Wake □off□, or Sleep □off□ values

These quality checks were performed on a total of 797 observations (of which only n=51 retrospectively logged). Eight entries (1%) failed Quality check #1, and 14 entries (1.76%) failed Quality check #2. These invalid entries were adjusted manually by referring to the raw files. We describe how these adjustments were applied in the **Supplementary Materials** (Data cleaning of Wear log entries: details on manually adjusted entries). After these adjustments, a rerun of the quality checks showed no erroneous entries after applying these corrections. The next step was to only include all entries up until and including the “Putting the light glasses on” entry after sleep on the last study day (Monday), before the participants returned to the laboratory to return the materials. Lastly, the Wear log entries were cleaned, arranged chronologically for each participant and integrated into the light logger dataset. This led to a final data frame where each light logger measurement, including a measurement of illuminance and activity, was labelled based on the Wear log *State* (Wake □on□, Wake □off□, Sleep □off□).

##### Data cleaning of button presses and low illuminance clusters

For the other two sources of non-wear information, namely the button presses and clusters of low illuminance levels, pre-processing was less extensive than for Wear log entries. Button presses within the same 60-second interval were considered invalid, i.e. caused by the participant accidentally pressing the button more than once to indicate the same (non-)wear action. These invalid consecutive button presses were removed, so only the first was kept. Regarding low illuminance clusters, the mEDI values corresponding to the Wear log Sleep □off□ state were excluded before implementing our custom-made algorithm, as the scope was to detect either wear or non-wear time during waking hours. Further transformations on illuminance levels throughout data analysis are detailed in the Results sections. When visualising illuminance levels on a logarithmic scale (**Figure 7A**), 0 values were handled by adding a value of 1 to all mEDI values. However, all calculations related to mEDI were performed without conversion to logarithmic scale and on the raw data, which included values of mEDI=0 lux.

#### Description of wear status based on ground truth

The first part of our analysis consists of a description of wear status based on the ground truth, i.e. the (non-)wear information obtained from the Wear log. Specifically, we examined the time spent in each Wear log state across the week, the distribution of Wake □off□ status during weekdays and weekend days, and the timing of Wake □off□ status across the day.

#### Analysis of button presses

To compare non-wear information obtained by the button presses to the ground truth, we examined whether the start and end of each Wake □off□ interval aligned with a button press. More specifically, we classified each Wake □off□ interval based on the presence of a button press within a given time window from the Wear log timestamp. Non-wear intervals were categorised as follows: intervals not bounded by a button press at the start and/or the end were labelled as open, while intervals bounded by two button presses, one at the start and one at the end, were labelled as closed. To investigate how the percentage of open and closed intervals varied with increasing time window lengths, we performed this analysis using eight windows ranging from one to eight minutes, increasing in one-minute steps.

#### Analysis of low illuminance clusters and low activity clusters

To compare non-wear information obtained by the use of the black bag to that of our ground truth, we used an algorithm (Hartmeyer et al., 2024) to detect clusters of low illuminance (melanopic equivalent daylight illuminance, mEDI) values in the light logger dataset. This algorithm takes three input parameters: (1) A maximum threshold for the boolean variable of interest, for which the clusters should be found (in our case, mEDI), (2) a minimum length of the cluster, and (3) a maximum interruption length of the cluster by values above threshold (in our case, mEDI values above threshold). With these three input parameters, the algorithm labels every illuminance observation in a participant’s light exposure dataset as belonging to a cluster of low illuminance levels or not. To assess the performance of this cluster detection algorithm in identifying the same non-wear intervals identified by our ground truth, we constructed a precision-recall (PR) curve. PR curves provide a graphical illustration of a classifier’s performance across several threshold values (in our case, mEDI values), with positive predictive value (PPV, or precision) on the Y-axis and true positive rate (TRP, or recall) on the X-axis. In the context of our classifier, precision indicates the number of instances that were predicted as non-wear by the algorithm and were non-wear according to the Wear log. On the other hand, recall indicates the number of instances correctly identified as non-wear by the algorithm out of all actual non-wear instances (i.e. Wear log Wake □off□ instances). Importantly, PR curves are preferred to commonly used receiver operating characteristic (ROC) curves in scenarios of class imbalance, as in our dataset. In this case, the small percentage of non-wear relative to wear intervals could risk overstating the algorithm’s performance, when evaluating it with a ROC curve (Cook & Ramadas, 2020).

To define the optimal combination of the minimum cluster length and maximum interruption length to detect clusters of low illuminance in our dataset, PR curves were first built with varying thresholds of these two variables, while keeping a threshold of <1 mEDI lux (Supplementary materials, **Figure S2 and Figure S4**). A minimum length of clusters of 21 minutes and a maximum interruption of 0 minutes led to the highest F1 score (F1=0.78). F1 is a metric representing the harmonic mean of precision and recall, with values ranging from 0 to 1 (Cook & Ramadas, 2020). Thus, these input parameters were used in our algorithm to build the PR curve with varying mEDI thresholds. Because clusters of low activity are often used to detect non-wear in field studies investigating sleep and light exposure with wearables similar to our light logger (Hartmeyer et al., 2022), we implemented our algorithm to also detect clusters of low activity in our dataset. Activity was quantified from the ActLumus built-in accelerometer as proportional integrating measure (PIM), a quantity directly calculated by the device throughout the study. As for illuminance, we first built PR curves for minimum cluster length and maximum cluster interruption length, while keeping an activity threshold of <5 (Supplementary materials, **Figure S3 and Figure S4**). A minimum cluster duration of 12 minutes, and a maximum cluster interruption of 0 minutes led to the highest F1 score (F1=0.52), and this combination was then used to build a PR curve with varying PIM thresholds.

#### Comparison of light exposure metrics

To determine whether handling non-wear intervals identified by either the Wear log or by the algorithm would result in different values of light exposure metrics compared to not handling non-wear intervals, we calculated 14 commonly used light exposure metrics (Hartmeyer & Andersen, 2023; Zauner, Hartmeyer, et al., 2025) and compared them across three variants:

1. Raw dataset: All mEDI values included, i.e., ignoring non-wear information;
2. Wear log-corrected dataset: All mEDI values corresponding to a *Wake off* interval replaced with “NA”, R’s standard placeholder for missing values;
3. Algorithm-corrected dataset: All mEDI values corresponding to algorithm-detected clusters of low illuminance replaced with “NA”.

Specifically, we calculated the following 14 metrics:

- Metrics representing temporal variability of light exposure: across days (interdaily stability, IS) and within days (intradaily variability, IV) (n=2);
- Metrics quantifying timing-related light exposure characteristics: Mean timing of light above 250 lux (MLiT_250_), mean timing of light above 1000 lux (MLiT_1000_), first timing of light above 10 lux (FLiT_10_), first timing of light above 250 lux (FLiT_250_), first timing of light above 1000 lux (FLiT_1000_), last timing of light above 10 lux (LLiT_10_), last timing of light above 250 lux (LLiT_250_), last timing of light above 1000 lux (LLiT_1000_) (n=8);
- Metrics quantifying duration-related light exposure characteristics: Time above 250 lux (TAT_250_), and time above 1000 lux (TAT_1000_) (n=2).
- Metrics quantifying intensity of light levels: average light exposure during daytime and mean across the brightest 10 hours (M10m).

All metrics were calculated for every day of the study where full light exposure data were available (n=6, since the first Monday was incomplete for all participants). IS and IV were calculated directly on the entire dataset, the outcome being a single participant value for each metric. Average light exposure during daytime was also calculated on the entire dataset, however, only for daytime periods, defined by the time from civil dawn until civil dusk, using photoperiod information at our specific latitude and longitude coordinates (see https://tscnlab.github.io/LightLogR/reference/photoperiod.html for more information). For the remaining metrics, values were first calculated for each of the six study days, and then mean and standard deviation values were calculated across these six values. This led to a single mean and standard deviation participant value for each metric. For average daytime light exposure and M10m, mEDI values were first log10 transformed, and a small value was added to 0 values prior to transformation. Values were then back-transformed (Zauner et al., 2025). Differences in the selected 14 metrics across the three datasets were assessed using Friedman test due non-normality of the data. Post-hoc pairwise comparisons were then performed using Wilcoxon test for paired data, and corrected for multiple comparisons with the Benjamini-Hochberg method (Benjamini & Hochberg, 1995). Results are presented here using a significance threshold of 0.05. Finally, to further explore differences in metrics across the three datasets beyond significance testing, we conducted equivalence testing on each post-hoc pairwise comparison to probe equivalence in light exposure metrics. Given the non-normal distribution of the data and the relatively small sample size, we employed bootstrapping to estimate the sampling distribution of the effect size for each pairwise comparison. To define the smallest effect size of interest (SESOI) for equivalence testing, we drew on prior literature examining weekday-to-weekend variation in personal light exposure (lux*hours), a common covariate in light exposure analysis. Based on large datasets from Switzerland (n=20 participants, 554 participant-days) and Malaysia (n=19 participants, 509 participant-days), where participants wore a light-measuring device continuously for one month (Biller, Zauner, et al., 2024), the observed effect sizes were 0.27 and 0.33, respectively. We therefore set the SESOI at *d*=0.3 (Cohen’s *d*, i.e., mean difference in metric divided by the standard deviation; Cohen, 2013). Equivalence was concluded when the SESOI (±0.3) completely covered 90% of the bootstrapped effect sizes (90% CI).

## Results

### Final dataset

A total of 30 participants enrolled in the study and began data collection. Of these, one participant withdrew from the study, and participation was terminated early for the other two participants. This was due to a lack of compliance with and understanding of the study procedure, whereby participants did not fill in the daily questionnaires, despite receiving various prompts from the researchers. This led to a sample of 27 participants who completed the study. One participant was excluded from the analyses because of technical problems with the light logger leading to unrecoverable data loss. Thus, our analyses were performed on a final sample of n=26 aged between 22 and 46 years (mean±1SD: 28±5 years, 14 female).

### Description of wear status based on ground truth

To explore how participants engaged in non-wear behaviours across the week, we first visually inspected light exposure, button presses and self-reported (Wear log) wear status for each participant across all study days (**Figure 4**). This allowed us to gain a qualitative understanding of how the three sources of non-wear (Wear log, button presses, and use of black bag) compared in our study. To quantify wear status according to our ground truth, we then calculated the amount of time participants spent in each Wear log state throughout the week: Wake ‘on□, Wake □off□ and Sleep □off□ (**Figure 5A**). Based on Wear log entries, participants spent 57.7% of their total participation time wearing the light logger (Wake □on□ mean, SD=6.3%, range= 42.2%–72.7%), 5.4% of time not wearing the light logger (Wake □off□ mean, SD=3.8%, range=0.3%–13.0%), and 37.0% of time with the light logger off during sleep (Sleep □off□ mean, SD=4.6%, range=25.9%–46.1%). When comparing non-wear time between weekdays and weekend days, a similar trend was observed, with participants spending a small amount of time not wearing the light glasses on weekdays (mean±SD= 4.7±3.8%, range=0%–16.0%) as well as during weekend days (mean±SD=7.0±8.0%, range=0%–29.9%) (**Figure 5B**). Moreover, we investigated non-wear time during wake periods only, as the participants were instructed to remove the light glasses while sleeping. Non-wear time across participants was 8.6±6.2% (mean±SD). To gain a better understanding of how the self-reported sleeping periods in the Wear log (Sleep ‘off’) related to participants’ actual sleep time, we compared these timestamps with participants’ self-reported sleep time measured with a daily sleep diary. The results highlighted that, for most participants, the two measures differed by less than 30 minutes (**Figure S5)**.

**Figure 4.**
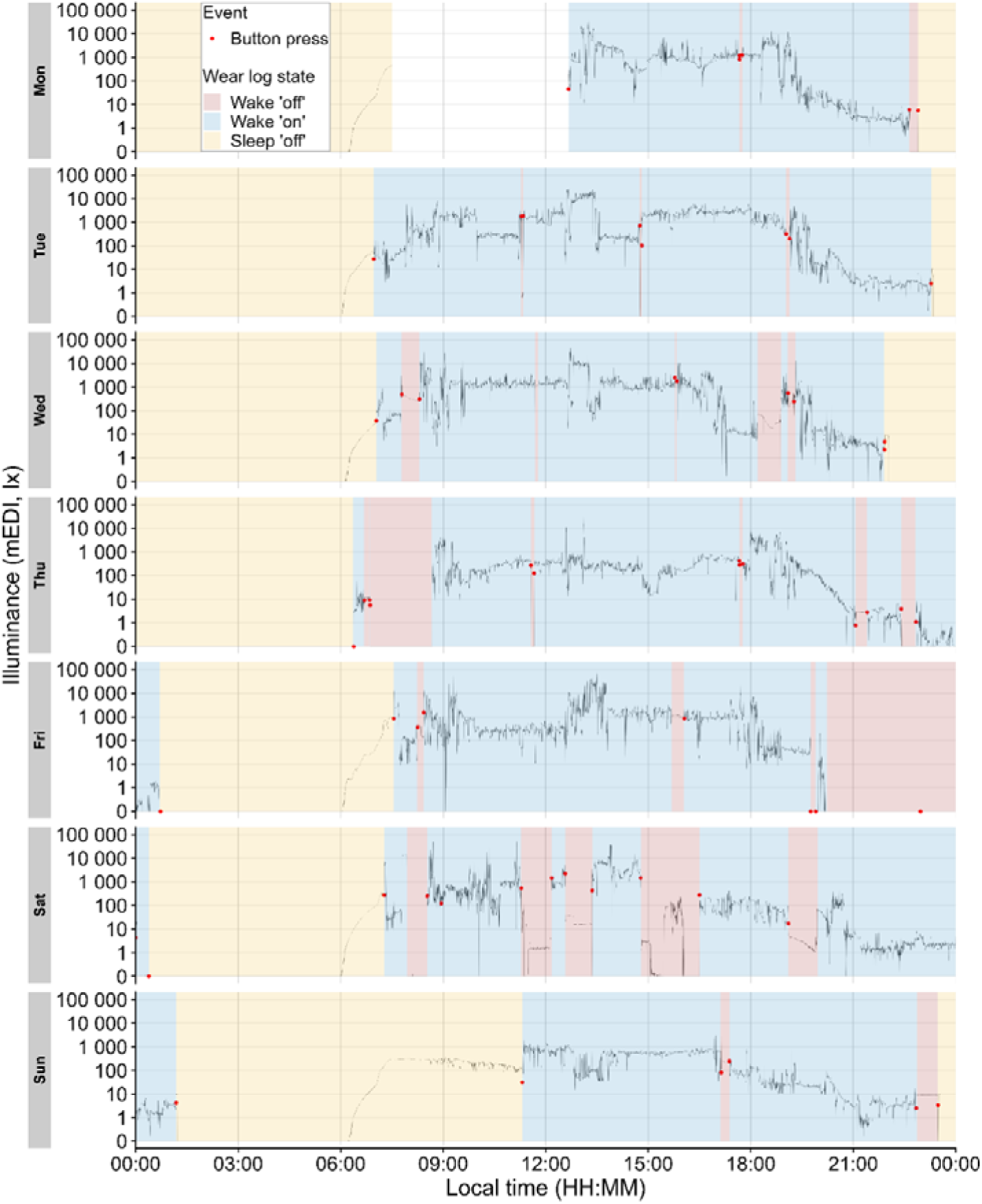
Exemplary light exposure and wear status for a participant (n=1) across the study. *Note.* The continuous black line shows mEDI as measured by the light glasses. Red points represent button presses performed by participants in case of any (non-)wear behaviour. Shaded areas represent self-reported wear status (Wear log).

**Figure 5.**
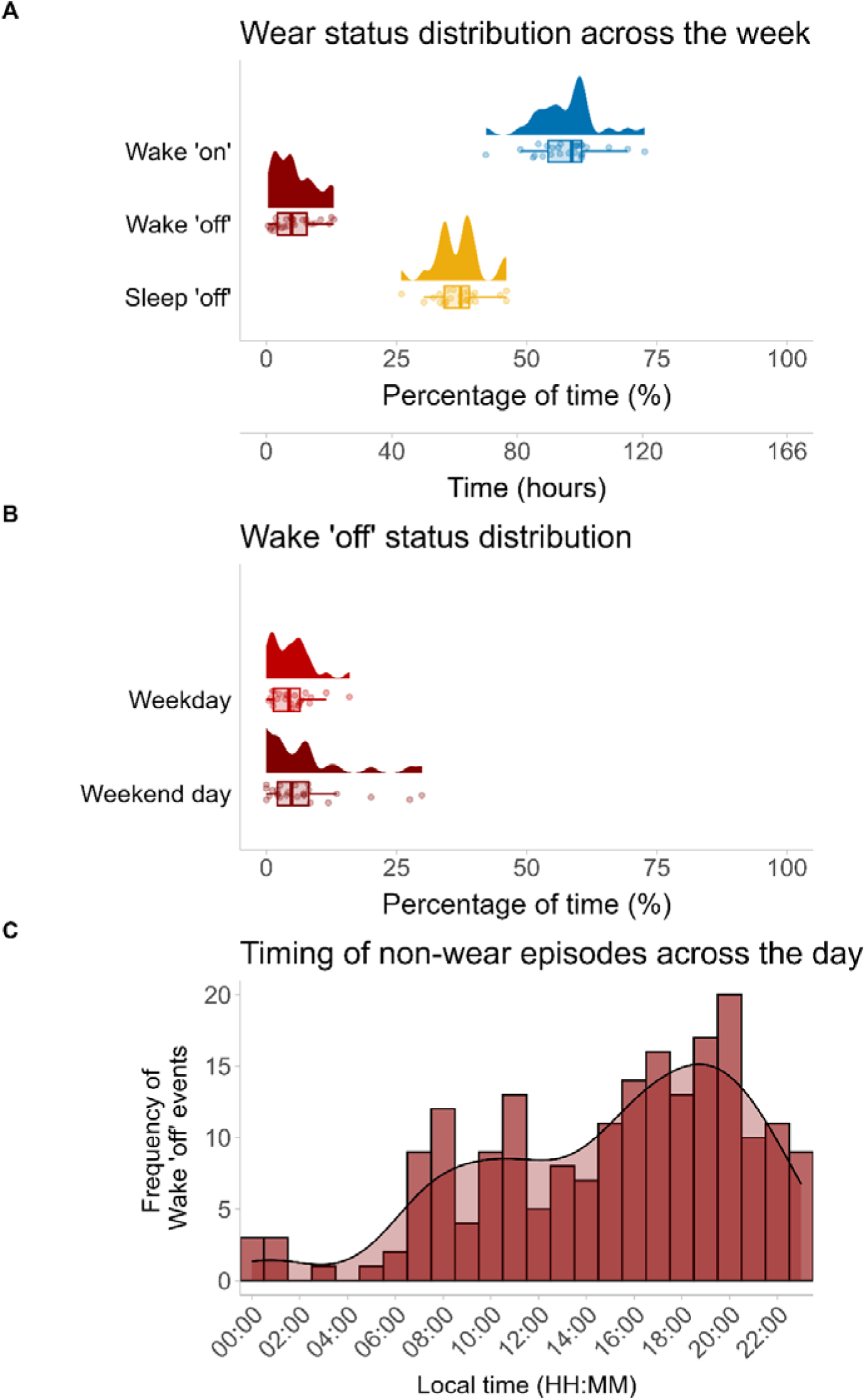
Distribution of self-reported wear status throughout the week and the day. *Note.* (A) Percentage of time spent in each Wear log state relative to individual study duration, and equivalent number of hours. Study duration slightly differed for each participant, as they all started the study at different times on the first Monday. (B) Percentage of time spent not wearing the light glasses (Wake ‘off’ state) relative to total weekday and weekend day time. (C) Frequency of non-wear episodes (Y-axis) over timing of non-wear episodes (X-axis). Timing of non-wear episodes (X-axis) was calculated as centre point of each Wake ‘off’ interval.

To further examine patterns in participants’ non-wear behaviour, we inspected duration and timing of reported non-wear episodes across the day. The median duration of non-wear was 00:37, with the 25^th^ and 75^th^ percentiles at 00:19 and 01:27, respectively. The minimum and maximum durations were 00:02 and 10:24 (all values reported as hh:mm). As a proxy of non-wear timing, we calculated the centre point of each Wake □off□ episode (**Figure 5C**). Non-wear episodes occurred continuously across 24 hours, with the exception of two time windows during night hours (between 02:00 and 03:00 and between 04:00 and 05:00). Density estimates for non-wear episodes showed an increase throughout the day and reached a peak at 20:00 with 20 non-wear episodes falling into this time bin. The number of non-wear time episodes decreased in the evening hours.

### Analysis of button presses

We next compared non-wear information obtained by the button presses to non-wear status information from our ground truth. Visual inspection suggested that only a few button presses (n=15) were not performed in proximity of a self-reported change in wear status. We examined the presence of button presses at the start and end of each Wake □off□ interval using different time windows. When considering button presses within a 1-minute window from a start and/or end of a Wake □off□ interval, only 64.1% of n=198 intervals were bounded by a button press on both ends and thus classified as closed. Expanding window sizes to more liberal cut-offs of 2, 3, 4, 5 and 6 minutes increased this number to 85.4%, 87.4%, 88.4%, 88.9%, and 89.9%, respectively. Adopting longer window sizes of 7 and 8 minutes did not increase the proportion of Wake □off□ intervals classified as closed, with the percentage of these intervals seemingly asymptotic at 88.9% for these cut-offs (**Figure 6**).

**Figure 6.**
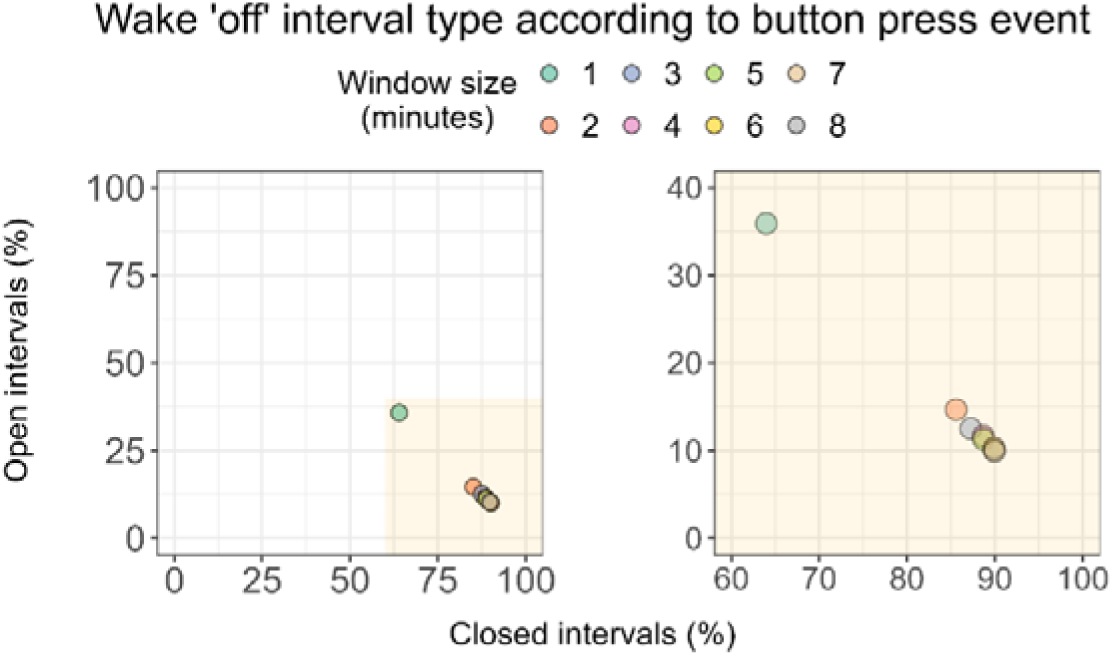
Classification of non-wear intervals based on presence of button press at the start and end of the interval. *Note.* Each Wear log Wake □off□ interval (n=198) was classified as open or closed based on the presence of a button press within a specified time window. Note that button presses at both the start and end of an interval were needed for an interval to qualify as closed.

### Algorithmic approach for non-wear detection

To explore the possibility of using clusters of low illuminance and low activity in our dataset as proxy of non-wear, we first examined illuminance and activity levels across Wear log states (**Figure 7A**). Participants’ median illuminance and median activity levels across the study showed a similar pattern, whereby similarly low mEDI and activity values were observed during Wake □off□ and Sleep □off□ intervals (mEDI mean±SD in Wake □off□ and Sleep □off□: 0.4±1.2 lux and 0.01±0.07 lux, respectively; activity mean±SD in Wake □off□ and Sleep □off□: 4.5±22.8 and 0.0±0.0, respectively). On the other hand, higher median illuminance and activity values were observed during Wake □on□ intervals (mean±SD: 62.4±66.6 lux and 61.5±46.1, respectively). Based on this observation, we used a previously developed algorithm (Hartmeyer et al., 2024) for detecting low illuminance clusters and low activity clusters in our dataset. The performance of this algorithm was examined using a precision-recall (PR) curve with varying maximum mEDI thresholds (from 1 lux to 10 lux, at 1-step increments), and activity thresholds (from 5 to 50, at increments of 5) (Figure 7B). Overall, detecting clusters of low illuminance performed better than detecting clusters of low activity in predicting non-wear. This difference was reflected in the F1 score, with the highest score for mEDI F1=0.78 (with a mEDI threshold of 1 lux) and the higher score for activity F1=0.52 (with an activity threshold of 5).

For illuminance, setting the threshold to smaller mEDI values for cluster detection led to a higher PPV (or precision) compared to higher mEDI thresholds (PPV=0.71 for <1 mEDI lux and PPV=0.33 for <10 mEDI lux). The gain in precision with decreasing mEDI threshold values came at a minimal cost for the TRP (or recall), with TRP=0.89 and TRP=0.81 for thresholds of <10 mEDI lux and <1 mEDI lux, respectively. Furthermore, we tested the algorithm performance separately for daytime periods (starting at dawn and ending at dusk) and nighttime periods (starting from dusk and ending at dawn). The PR curve revealed a better performance during daytime periods (highest F1=0.80, with mEDI threshold of 2 lux), compared to nighttime periods (highest F1=0.76, with mEDI threshold of 1 lux) (**Figure S6**). Of note, daytime performance consistently exceeded nighttime performance, with the lowest daytime performance (F1=0.68 for mEDI threshold <10 lux) still higher than nighttime performance for all mEDI thresholds except <1 lux and <2 lux. As the PR curve in **Figure 7B** shows the aggregate classification accuracy, we next probed whether participants had idiosyncratic non-wear behaviours and analysed if classification accuracy depended on each participant. This was indeed the case (**Figure 8**). Furthermore, we did not identify a clear pattern for classification accuracy of low illuminance clusters and participants’ self-reported compliance with using the black bag during non-wear time. For example, for a participant who reportedly used the black bag for all non-wear (Wake □off□ events), both precision and recall of the classifier were low (**Figure 8, PID 214**). Likewise, for a participant who reportedly never placed the light glasses in the black bag when taking them off, the recall of the classifier was very high (**Figure 8, PID 218**).

**Figure 7.**
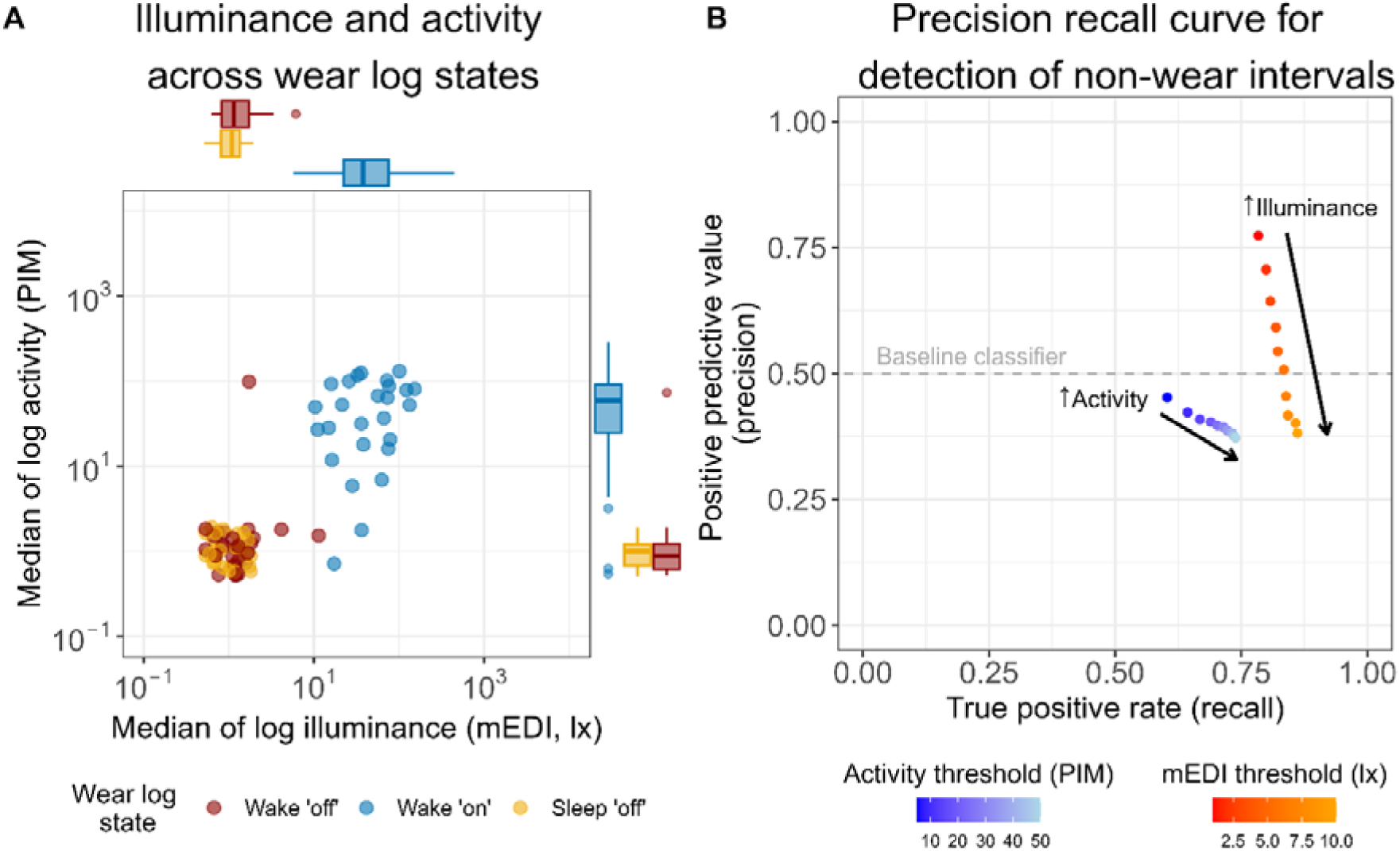
Classification of non-wear time based on detection of clusters of low illuminance and low activity. *Note.* (A) Median values of log activity and illuminance throughout the week based on Wear log state. Every data point indicates the weekly median value for a participant (n=26 points for each Wear log state). Note that for this visualisation, a value of one was added to all mEDI and PIM raw values prior to transformation to logarithmic scale. (B) Precision recall curve for detection of non-wear intervals based on clusters low illuminance (red-orange colour scale) and low activity (blue-light blue colour scale), as detected via our algorithm. A minimum cluster length of 21 minutes and a maximum interruption length of 0 minutes were used as input parameters in our algorithm for illuminance. For activity, a minimum cluster length of 12 minutes and a maximum interruption length of 0 minutes were used. Each data point on the curve represents a maximum threshold (mEDI: 1 lux to 10 lux, at 1-step increments; PIM: 5 to 50, at 5-step increments).

**Figure 8.**
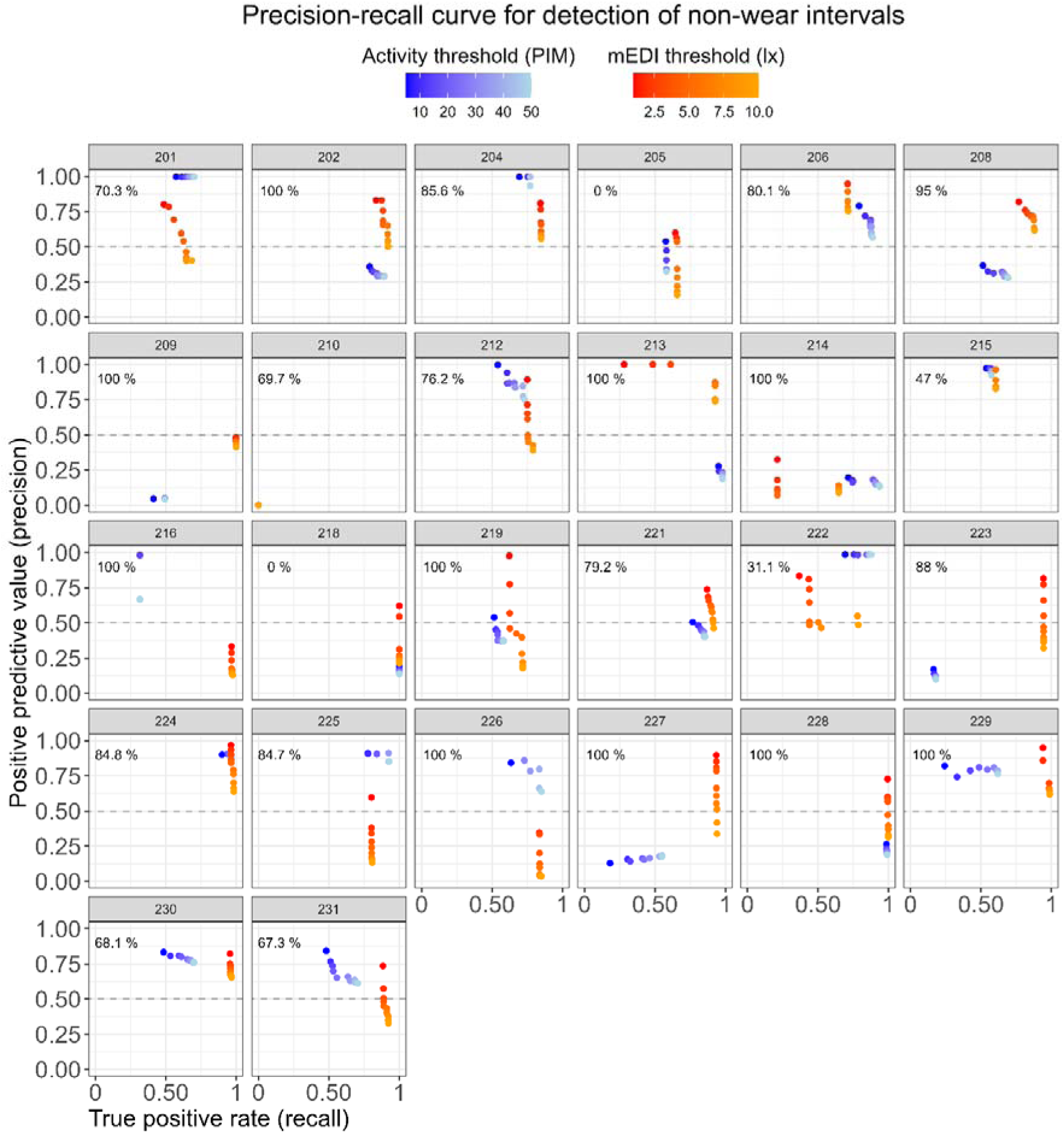
Individual precision-recall curves for detection of non-wear intervals based on clusters of low illuminance and low activity, as calculated by the algorithm. *Note.* A minimum cluster length of 21 minutes and a maximum interruption length of 0 minutes were used as input parameters in our algorithm for illuminance (red-orange colour scale). For activity, a minimum cluster length of 12 minutes and a maximum interruption length of 0 minutes were used (blue-light blue colour scale). Each panel represents a participant, and each data point within a panel represents a maximum threshold (mEDI: 1 lux to 10 lux, at 1-step increments; PIM: 5 to 50, at 5-step increments). The percentage value displayed on the top left of each panel shows the frequency of bag use relative to the participant’s self-reported non-wear episodes.

#### Examination of algorithm misclassification

Since detecting clusters of low activity would be an easy-to-implement strategy to detect non-wear, we examined reasons behind the poor performance of the algorithm for detection of non-wear based on clusters of low activity. We first analysed whether using PIM as activity parameter was a justified choice, and whether changing activity parameter would yield a different algorithm performance. ActLumus files contain three parameters that quantify activity: proportional integration mode (PIM, a measure of activity levels), time above threshold (TAT, a measure of time spent in motion), and zero crossing mode (ZCM, a measure of frequency of movement) (Pilz et al., 2022). Our results indicate that the performance of our algorithm was similarly poor for all three activity-quantifying parameters (Supplementary materials, **Figure S7**). Furthermore, we investigated whether pre-processing activity (PIM) values would lead to a better classification performance. We applied several pre-processing methods including log-transformation, smoothing, and normalisation of PIM values, and showed that none of these strategies benefitted classification performance (Supplementary materials, **Figure S8**). Given the superior performance in detecting clusters of low illuminance using a mEDI threshold of <1 lux (F1=0.78), we then applied our algorithm to the dataset using this threshold, and focused subsequent analyses on illuminance rather than activity. In order to understand in which circumstances our algorithm failed at predicting non-wear (resulting in false positive and false negative instances), we next analysed characteristics of these instances that might have contributed to misclassification. Specifically, we calculated the duration of each misclassified instance, the mean melanopic illuminance during this interval, and whether the participants reported using the bag during this time. Using these three variables, we identified four potential causes of misclassification and labelled each false negative and false positive interval in our dataset accordingly:

1. Algorithm limitation or bag not actually used: False negative intervals (>3 minutes) where participants reported using the bag and with mean mEDI >1 lux. These false negatives could be caused by participants not using the black bag despite reporting doing so, or by the algorithm input parameters preventing correct classification (e.g. cluster duration <21 minutes).
2. Bag not used: False negative intervals of any duration where participants reported not using the bag and with mean mEDI >1 lux. In this case, since the bag was reportedly not used and the mEDI values suggest similarly, no clusters of low illuminance could be detected by the algorithm.
3. Low illuminance during wear: False positive intervals (<3 minutes) with mean mEDI ≤1 lux. In this case, individuals received low illuminance during wear time, leading to the algorithm detecting these intervals as non-wear clusters.
4. Transition state: False negative intervals (≤3 minutes) where participants reported using the bag and with mean mEDI >1 lux, and false positive intervals (≤3 minutes) with mean mEDI ≤1 lux. These brief intervals likely represent transitions from one Wear log state to the next one, where a short time lag between logging wear status on the Wear log and placement or removal of the light glasses from the black bag can lead to misclassification.

A summary of misclassified and correctly classified instances can be found in **Table 1**. Furthermore, examples of the algorithm misclassification instances in **Table 1** are shown in an exemplary participant (**Figure 9**). Due to the high number of transition states identified as potential causes of our algorithm’s misclassification (n=210, or 21.5% of 978 total instances), we implemented a modification to the algorithm-detected clusters of low illuminance. Specifically, for each cluster of low illuminance values (i.e. a non-wear cluster), we checked for a transition state label, identified as described above, immediately before or immediately after the cluster. If a transition state was detected, the non-wear cluster was expanded as follows. In case of a transition state detected before the cluster, all observations from the start of the transition state to the start of the non-wear cluster were re-labelled as non-wear. In the case of a transition state after the cluster, all observations from the end of the non-wear cluster to the end of the transition state were re-labelled as non-wear. This adjustment ensured that each low illuminance cluster adjacent to a transition state was “padded” with non-wear observations at both ends.

**Figure 9.**
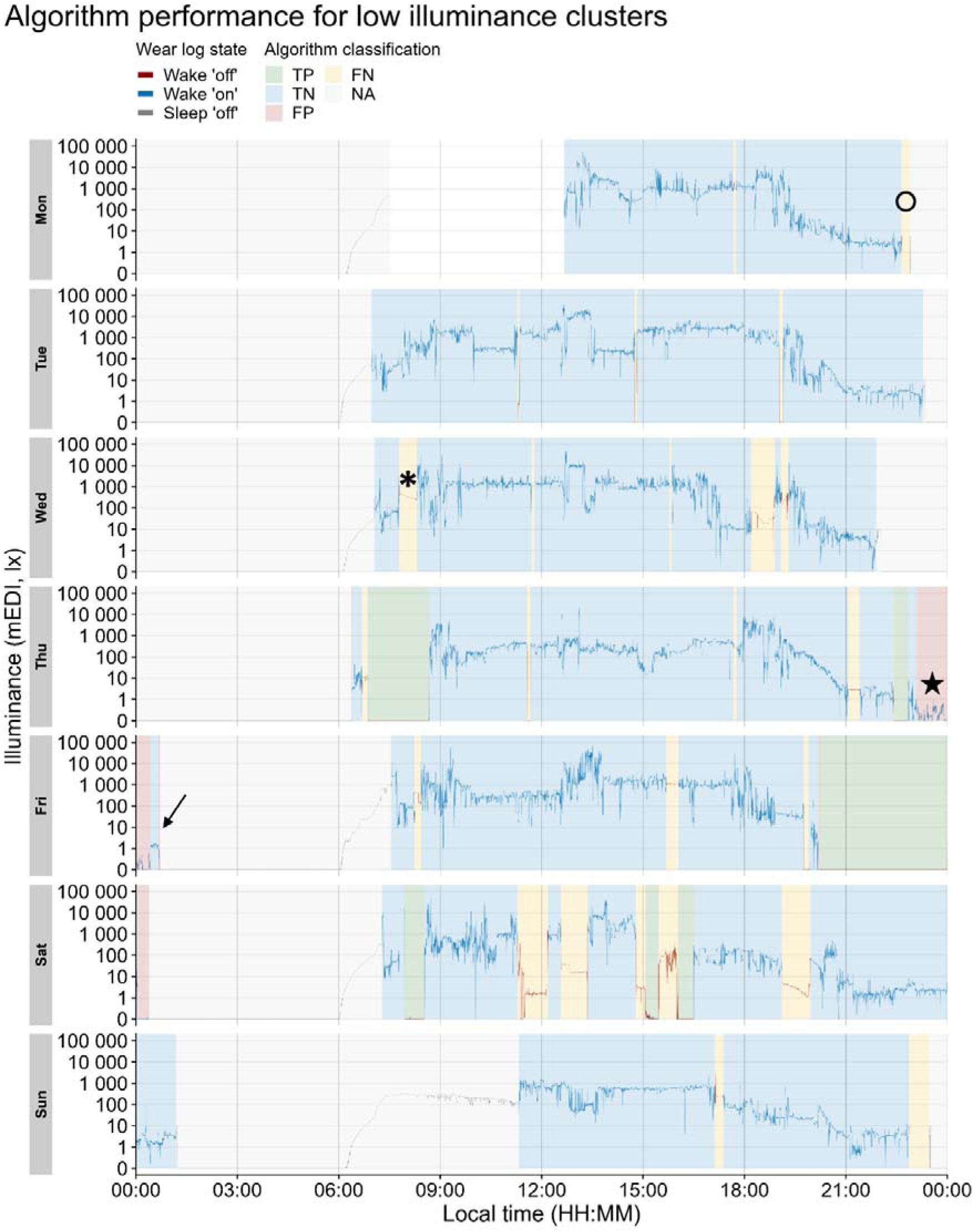
Example of misclassification instances for a participant (n=1) across the study. *Note.* Blue and green shaded areas represent correctly classified intervals (true negative and true positive, respectively). Yellow and red areas represent misclassified intervals (false negative and false positive, respectively). Grey shaded areas represent sleep episodes (excluded from the algorithm classification). The colour of the line representing light levels (mEDI) mirrors the wear status according to the ground truth (Wear log). The black symbols indicate examples of misclassified intervals as labelled in Table 1. Arrow: example of “Misclassified: Transition state”; Open circle: example of “Misclassified: Algorithm limitation or bag not actually used” instance; Asterisk: “Misclassified: Bag not used” instance; Star: “Misclassified: Low illuminance during wear” instance.

**Table 1.**
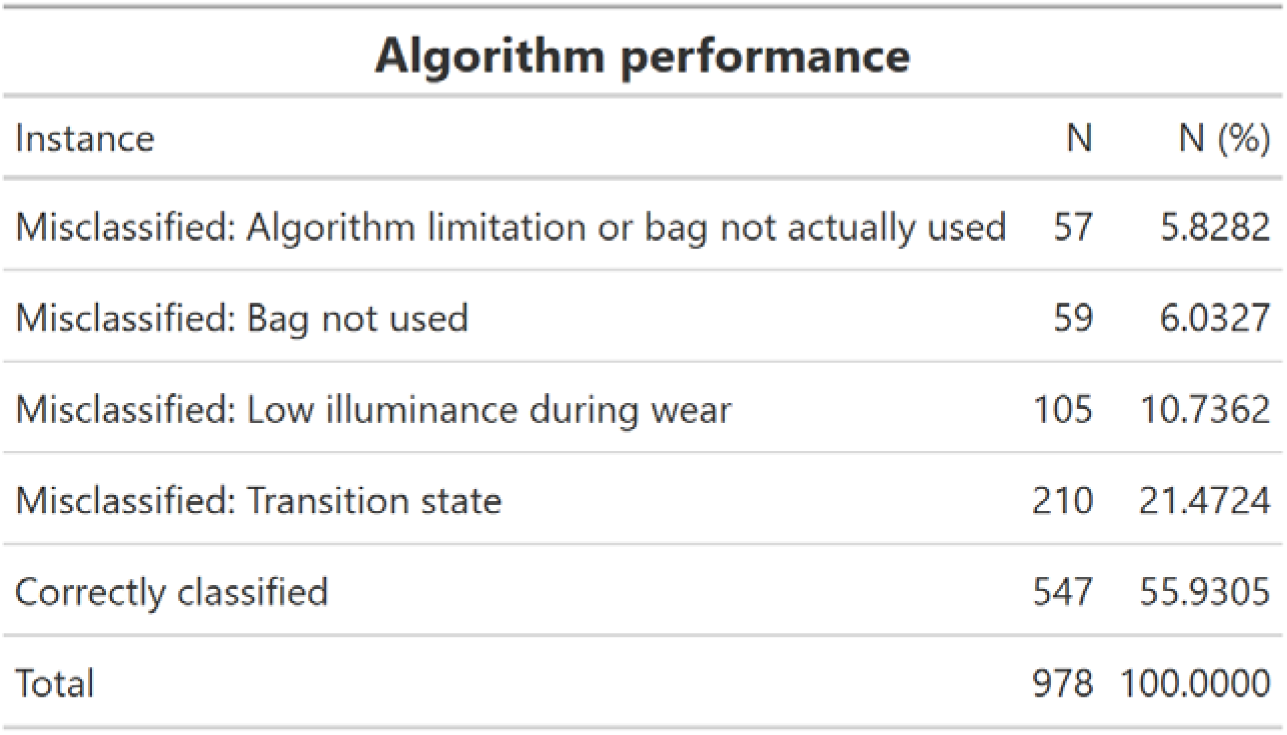
Summary of correctly classified and misclassified (non-)wear instances by the algorithm detecting low illuminance clusters. *Note.* Correctly classified instances represent true negative and true positive instances, and misclassified instances represent false positive and false negative instances. Potential reasons for misclassified instances were deducted from the duration of the misclassified instance, mean mEDI during this interval, and self-reported use of the black bag during this instance.

| Algorithm performance |  |  |
| --- | --- | --- |
| Instance | N | N (%) |
| Misclassified: Algorithm limitation or bag not actually used | 57 | 5.8282 |
| Misclassified: Bag not used | 59 | 6.0327 |
| Misclassified: Low illuminance during wear | 105 | 10.7362 |
| Misclassified: Transition state | 210 | 21.4724 |
| Correctly classified | 547 | 55.9305 |
| Total | 978 | 100.0000 |

### Comparison of light exposure metrics

To assess whether removal of non-wear time would affect light exposure metrics in our data, we calculated 14 light exposure metrics across three datasets.

The choice of light metrics was based on commonly used metrics quantifying timing-related characteristics (MLiT_250_, MLiT_1000_, FLiT_10_, FLiT_250_, FLiT_1000_, LLiT_10_, LLiT_250_, LLiT_1000_), duration-related characteristics (TAT_250_ and TAT_1000_), temporal dynamics (IS and IV) of light exposure patterns, as well as light levels (average light level during daytime hours and mean across the brightest 10 hours, M10m) (Hartmeyer & Andersen, 2023; Zauner et al., 2025). The relationship between each metric calculated in the raw dataset and the corrected, cleaned datasets (Wear log-corrected and algorithm-corrected), as well as a histogram of the delta of the metrics, can be found in **Figure 10**. To test for differences in metrics across the three derived datasets, we performed a Friedman test. The results indicated statistical differences in metrics between the three datasets from all four metric domains: LLiT_10_, LLiT_1000_, TAT_250_, TAT_1000_, average light exposure during daytime hours, M10m, and IV. No statistical difference was found for the following metrics quantifying timing-related patterns of light exposure and its temporal variability: MLiT_250_, MLiT_1000_, LLiT_250_, FLiT_10_, FLiT_250_, FLiT_1000_, and IS (**Table S1**). When examining post-hoc comparisons (n=21) using Wilcoxon signed-rank tests with Benjamini-Hochberg correction for multiple comparisons, n=9 pairwise comparisons reached statistical significance, corresponding to light exposure metrics related to duration-related patterns of light exposure as well as its intensity (TAT_250_, TAT_1000_, average daytime mEDI, and M10m). For metrics quantifying timing-related patterns of light exposure and its temporal variability that resulted in significantly different outcomes in the three-way comparison (LLiT_10_, LLiT_100_, and IV), no statistical difference across the pairwise comparisons was found. A full list of pairwise comparisons and adjusted p-values is provided in **Table 2**. Furthermore, the means and standard deviation values for all 14 metrics are reported in **Table S2**.

**Figure 10.**
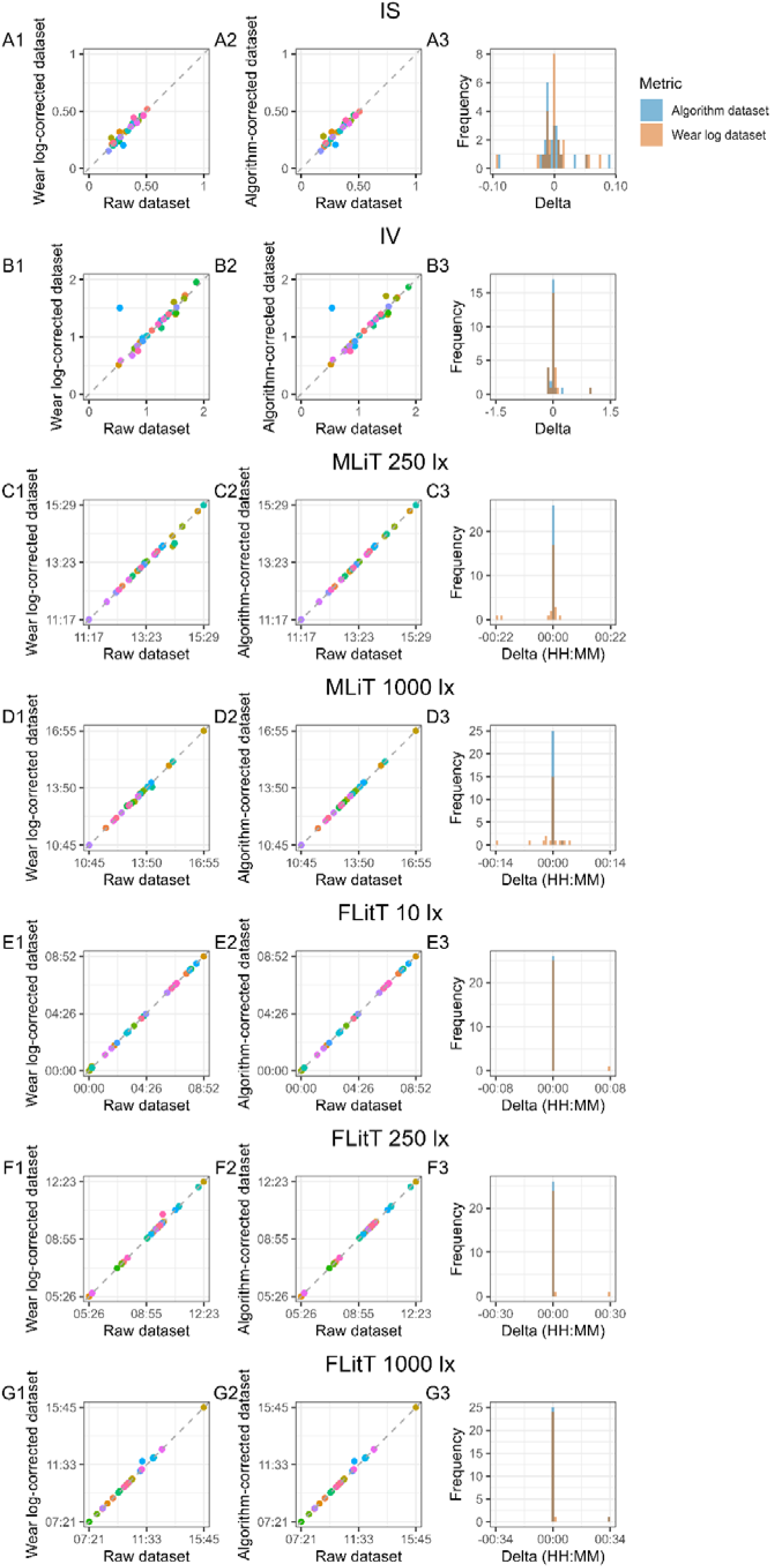
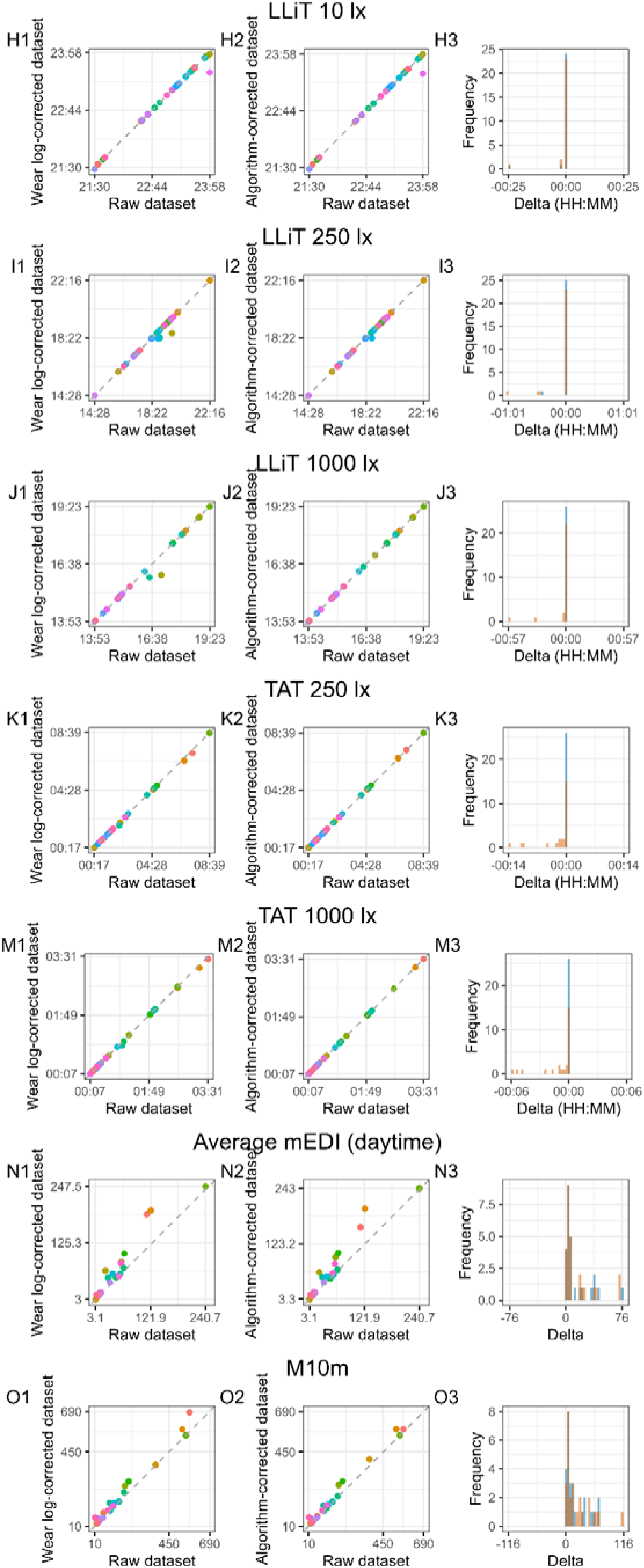
Comparison of 14 light exposure metrics (A-M) between the raw dataset and the clean (Wear log) dataset and between the raw dataset and the clean (algorithm) dataset. *Note.* For each metric, the first scatter plot shows the relationship between a given metric in the raw dataset and in the clean (Wear log) dataset (A1-O1), and a second scatterplot shows the relationship between a given metric in the raw dataset and in the clean (algorithm) dataset (A2-O2). Each data point in the scatterplot represents the value of the metric for a given participant (n=26). Distribution of the delta values for the clean (Wear log) dataset and clean (algorithm) dataset relative of the raw dataset are shown in A3-M3. Note that for timing metrics (C3-J3; MLiT^250^, MLiT^1000^, FLiT^10^, FLiT^250^, FLIT^1000^, LLiT^10^, LLiT^250^, LLiT^1000^), the label HH:MM on the X-axis of the histogram of the deltas represents local time, whereas for duration metrics (K3-M3; TAT^250^ and TAT^1000^), the HH:MM represents a duration. For the intensity metrics (average mEDI and M10m), the values shown in the graphs represent mEDI lux (N1-O1 and N2-O2).

**Table 2.**
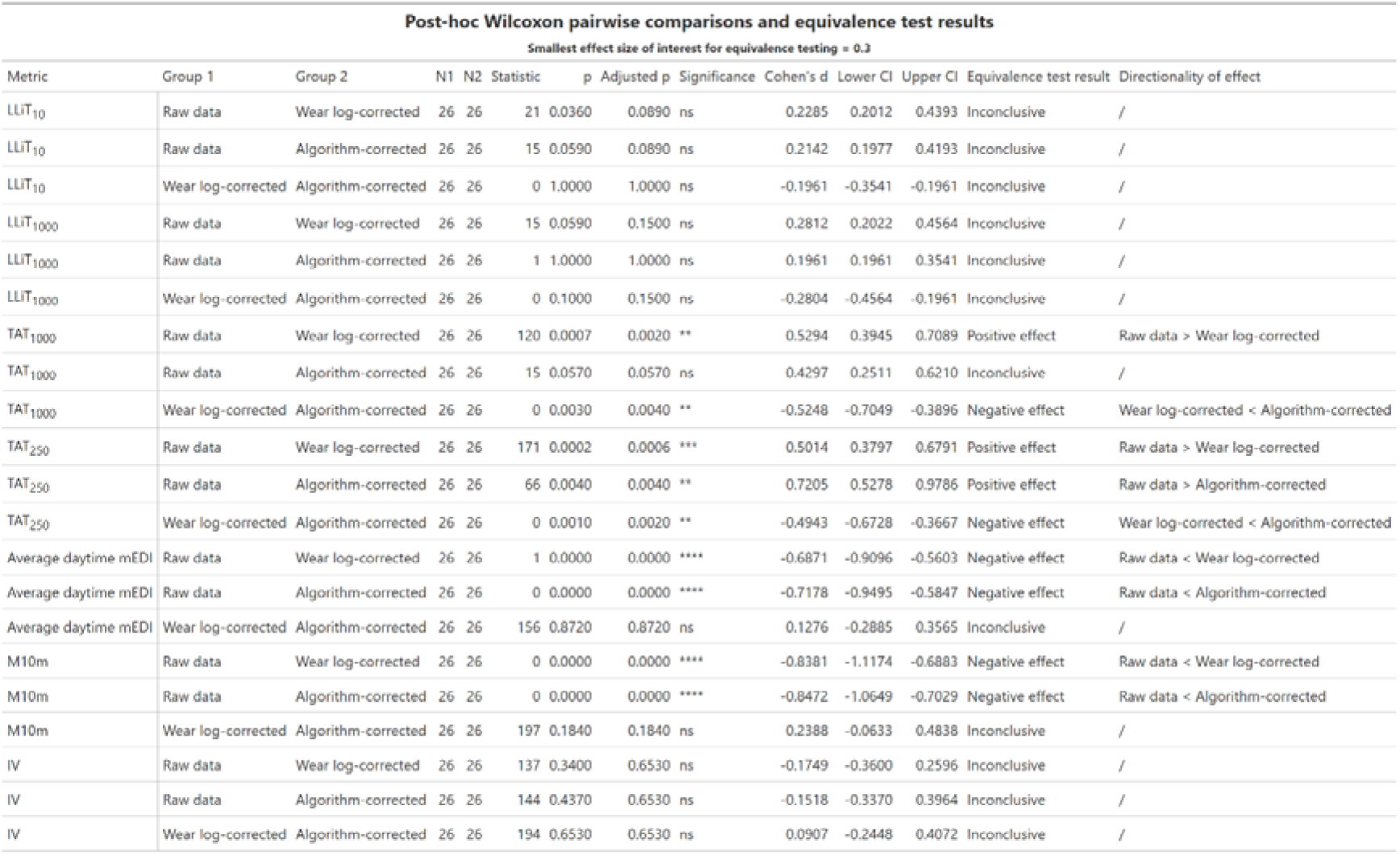
Post-hoc Wilcoxon signed rank test comparisons results and corresponding equivalence test results. *Note.* Post-hoc Wilcoxon signed rank tests were performed for all metrics where the Friedman test returned a significant difference. P-value adjustment was performed using the Benjamini-Hochberg method for multiple comparison correction. Significant values are reported at threshold p<0.05. Equivalence tests were also run on all post-hoc pairwise comparisons using a smallest effect size of interest (SESOI) of *d* = 0.3. For comparisons yielding to positive or negative effects in equivalence testing (i.e. both 90% confidence intervals entirely above or entirely below the SESOI), the directionality indicates which dataset had higher or lower values.

Among the 21 pairwise comparisons investigated, none were conclusively equivalent. For n=9 comparisons, a directed effect was found; i.e. the bootstrapped 90% confidence intervals of the effect size between metrics lay entirely above or below the SESOI region. Specifically, for metric quantifying duration-related patterns (TAT_250_ and TAT_1000_), raw dataset values were higher than Wear log-corrected values (TAT_1000_) as well as algorithm-corrected values (TAT_250_). The opposite trend was observed for metrics quantifying light intensity (average daytime mEDI and M10m): raw dataset values resulted lower than Wear log- and algorithm-corrected values. The remaining n=12 comparisons were inconclusive. Equivalence test results for each pairwise comparison are reported in **Table 2**, and corresponding bootstrapped effect size distributions are shown in **Figure S9**. A summary of differences in metrics from their uncorrected values (raw dataset), as well as for differences between correction methods (Wear log-corrected and algorithm-corrected), can be found in **Table 3**. Furthermore, **while** our choice of SESOI for equivalence testing was based on literature for weekday-to-weekend effects, other SESOI thresholds can be relevant in other contexts. We thus further explored how different SESOI thresholds (0 to 1, at 0.1 steps) might influence the outcome of the equivalence tests. **Figure S10** illustrates the proportion of bootstrapped effect sizes falling outside of the SESOI region as a function of SESOI threshold, while **Figure S11** summarises how the proportion of equivalence conclusions changes depending on SESOI selection.

**Table 3.** Summary of positive, negative, and inconclusive effects identified via equivalence testing. *Note.* Summary of light exposure metrics that yielded statistically meaningful positive, negative or inconclusive effects in pairwise comparisons performed in equivalence testing (SESOI = ±0.3). The metrics are categorised by differences from the raw dataset (i.e. no correction; table columns) and differences between correction methods (i.e. Wear log-corrected and algorithm-corrected; table rows).

|  |  | Difference from no correction |  |
| --- | --- | --- | --- |
|  |  | Inconclusive | Yes |
| Difference between correction methods | Inconclusive | LLiT<sub>10</sub><br>LLiT<sub>1000</sub><br>IV<br>IS | Average daytime mEDI<br>M10m |
|  | Yes | / | TAT<sub>1000</sub><br>TAT<sub>250</sub> |

Uncorrected metric value is lower
Uncorrected metric value is higher

## Discussion

To the best of our knowledge, this is the first study to systematically assess the impact of non-wear intervals on light exposure metrics in ambulatory light logging studies. Our protocol, characterised by continuous data quality checks, was designed to yield a high-quality ground truth for non-wear intervals, i.e. the self-reported Wear log entries, essential to evaluate and compare different non-wear detection strategies. Our results indicated that button presses did not consistently mark the start and end of non-wear intervals, and that algorithmically detecting clusters of low illuminance resulting from black bag use had a reasonably strong performance against ground truth labels (F1=0.78). Lastly, we found that addressing non-wear intervals in our small, high-compliance dataset only had a marginal impact on some of the commonly used light exposure metrics examined.

### Description of wear status based on ground truth

As part of data pre-processing, we first visually inspected all three sources of non-wear. Relying on visual inspection alone to detect non-wear intervals is time consuming and requires training (Pilz et al., 2022), but it can be used as an initial step for exploring raw data. Visual inspection allowed us to gain a better understanding of participant compliance and concordance between non-wear sources, and provided a qualitative insight into which automated detections (i.e., non-wear algorithms) could be applied to our data. Visualisation of raw light exposure data is possible using open-access packages in programming environments such as the *LightLogR* package (Zauner et al., 2025) or the *pyLight* submodule of pyActigraphy (Hammad et al., 2024), the former also offering options to visualise multiple non-wear sources in the same graph, as shown in Figure 4 of this paper.

When examining self-reported Wear log entries, the relatively low non-wear time (mean±SD=5.4±3.8% of total participation time and 8.6±6.2% of wake-only time) across the sample indicates that participants were compliant with study instructions, spending a substantial amount of time wearing the light glasses during their daily activities. Compliance rates were high for both weekdays and weekend days. Our results align closely with the findings of a recent study where participants wore a near-corneal plane light logger for five days, a duration similar to that of our study. The authors report that, despite participants documenting discomfort and unwanted attention in public settings when wearing the device, adherence to the study instructions was high (89% during waking hours, excluding sleep time) (Stefani et al., 2024). As to the timing of non-wear, the highest number of reported non-wear episodes in our dataset coincided with early evening hours, peaking at 20:00. Stefani and colleagues (2024) also reported slightly higher wear compliance during daytime compared to evening hours during weekdays (92% of recording time during daytime, 71% of recording time the evening). While we lack additional data to understand which specific situation prompted the removal of the light glasses at this time, we speculate that reasons for removal might be similar to those reported in other studies that used near-corneal level light loggers: being in the shower, being in public spaces where wearing the light glasses is thought to cause undesirable reactions from others, performing sports, or experiencing discomfort (Balajadia et al., 2023; Stefani et al., 2024).

### Analysis of button presses

When turning to button presses marking the start or end of a non-wear interval, our results indicated that button press events mostly happened within 1 minute to 6 minutes from the corresponding Wear log entry. Extending the time window beyond this range did not result in more non-wear intervals labelled as closed, suggesting that participants occasionally forgot to press the button to mark a non-wear event. To the best of our knowledge, no previous study has examined the concordance of button presses with self-reported non-wear time, although some studies mention instructing participants to use this when removing the light logger (Van der Maren et al., 2018), as well as using it in visual inspection of the data (Stefani et al., 2024). Our results suggest that button presses alone are unreliable for collecting accurate information on non-wear intervals. However, since several available light loggers have a built-in event marker (van Duijnhoven et al., 2024), collecting non-wear information using button presses does not add a significant burden to participants and can help validate non-wear self-reports. This strategy could especially benefit research studies in which non-wear self-reports are not as clean as in the current dataset, and no other source of non-wear is collected. Based on our findings, we recommend using cut-off criteria larger than 1 minute for automatic detection of button presses corresponding to a self-reported non-wear event, as stricter criteria could compromise non-wear validation.

### Data-driven non-wear detection

As part of our non-wear detection analysis, we applied an algorithm detecting clusters of low illuminance values caused by the use of the black bag during non-wear time, as well as clusters of low activity, which might also be indicative of non-wear time. Evaluating the performance of our algorithm using PR curves, our results indicate that collecting non-wear information by asking participants to place the light logger in a black bag might be an effective strategy to facilitate automated detection of non-wear time in ambulatory light exposure studies. When varying the input parameters (maximum mEDI levels, minimum cluster length, and maximum cluster interruption), we found that both the maximum mEDI threshold and the minimum cluster length had a clear influence on performance, whereas the maximum cluster interruption length had only a minor effect. Based on these findings, we recommend adopting a conservative threshold for maximum light levels (<1 mEDI lux). For the minimum cluster length, durations between 10 and 30 minutes yielded comparable F1 scores (0.76–0.78), likely reflecting the variable duration of non-wear episodes among participants. Therefore, in field studies where non-wear duration is uncontrolled, we suggest prioritising medium-length minimum cluster lengths (between 10 and 30 minutes) over very short ones (<10 minutes) to improve detection. More broadly, our findings highlight that maximum light level thresholds and minimum cluster length should be explicitly considered when developing more complex algorithms for non-wear detection to ensure both accuracy and generalizability across study designs.

Our results also highlight that photoperiod information (day and night) must be considered when deciding on an illuminance threshold for automated non-wear detection. When only looking at daytime periods (dawn to dusk), the strong contrast between environmental light (e.g. daylight or electric light) illuminance levels and the low black bag illuminance levels might lead to better algorithm performance. On the other hand, during the night period (dusk to dawn), the low intensity of the environmental light might impair the ability of the algorithm to accurately detect clusters of low illuminance due to black bag use. While for both daytime and nighttime lower mEDI thresholds led to better algorithm performance, higher mEDI thresholds during daytime (<9 and <10 lux) led to a similar performance to low mEDI thresholds during nighttime (<2 lux). Overall, this result suggests that when using an automated low-illuminance detection algorithm in periods of naturally lower light levels (i.e. nighttime), a more conservative illuminance threshold should be adopted. Considering the poorer performance of detecting low activity clusters compared to low illuminance clusters, it is important to consider that this was not always observed when looking at individual PR curves (**Figure 8**). Specifically, for some individuals, using low illuminance clusters to detect non-wear led to perfect recall, but using clusters of low activity led to greater precision. Furthermore, the observation that compliance with bag use did not seem to underlie better algorithm performance for illuminance might be explained by individuals placing the device in a dark space, such as a drawer, even when they did not place it in the black bag. We suggest that relying on both illuminance and activity (i.e., multi-modal input to our algorithm) could improve automated non-wear detection, particularly in accounting for non-compliance with black bag use. This approach would increase classification performance for non-wear intervals where the signal from the two variables is opposite, such as when the device sits on an illuminated surface (i.e., high light levels, low activity levels).

Upon examining the reasons behind our algorithm’s misclassifications, transition states were a major contributor to false positives and false negatives. We attribute this to a time mismatch in non-wear logging: While participants were instructed to first place the light glasses in the black bag and then log non-wear in the Wear log, they might have performed these actions in reverse order. As a result, the Wear log would indicate a non-wear period at time X, while the algorithm would detect a cluster of low illuminance only at time X+1 minute, when the light glasses were placed in the black bag. Consequently, the algorithm would label the instances between X and X+1 minute as false negatives. A similar issue occurs when putting the glasses back on after non-wear. If participants log this action before taking the light glasses out of the bag, the algorithm detection will result in false positive instances until the device is actually outside the bag and being worn. We addressed this transition state problem post-hoc by padding each non-wear cluster where a misclassification related to a transition state was identified. A further improvement to our algorithm would be to identify transition states dynamically as part of the classification process. It is important to note that our algorithmic approach might not be the only one suitable for identifying non-wear time in ambulatory light exposure studies. For example, since our algorithm employs a minimum length during which light or activity must be below a threshold value, short intervals are likely misclassified. Algorithms that do not rely on an interval but rather explore signal characteristics immediately preceding or following a non-wear interval might lead to better non-wear classification (Syed et al., 2021).

### Comparison of light exposure metrics

Our analysis on commonly used light exposure metrics revealed that accounting for non-wear time intervals only affected n=7 out of n=14 light metrics investigated. Of note, post-hoc tests revealed that for three (LLiT_10_, LLiT_100_, and IV) of the n=7 affected metrics, no statistical difference across pairwise comparisons was found. Furthermore, equivalence testing was inconclusive for a small effect size (SESOI=±0.3), indicating insufficient evidence that the differences in metrics were small enough to be considered practically equivalent. Thus, while the metrics are not significantly different between the methods of non-wear handling, they might be in more sensitive cases (i.e. with higher statistical power).

For n=9 out of n=21 post-hoc pairwise comparisons, a significant difference was found and equivalence testing indicated a directed effect. Importantly, with the exception of one metric type (TAT), equivalence testing indicated that differences were found only between the raw dataset and a corrected dataset, not between the Wear log-corrected set and the algorithm-corrected set. Interestingly, raw dataset values were higher than Wear log-corrected values for duration-related metrics, and the opposite trend was observed for metrics quantifying light intensity. These diverging trends might be due to duration metrics being sensitive to peaks of high light exposure (i.e. non-wear periods during which participants are not using the black bag, since these metrics calculate any value above threshold), and less sensitive to periods of non-wear in a dim or dark environment (i.e. non-wear periods during which the black bag is used, since these contribute to zero time above threshold). On the other hand, intensity-related metrics are more sensitive than duration metrics to whether or not low light observations in the data are filtered (i.e. low illuminance values resulting from black bag use). However, it is important to note that the differences in daily mean TAT_250_ and TAT_1000_ values between the raw and Wear log-corrected dataset are very small (∼2 minutes and ∼1 minute, respectively). This means that, despite the difference being significant, the magnitude of this difference is very small and likely driven by low variability in participant-level differences rather than a meaningful change in the metric. The lack of differences between light metrics could be explained by the timing of non-wear observed in our dataset. Our participants mostly reported removing the light glasses in the evening hours, when light levels tend to be low. Thus, the missing data during this period do not impact light metrics quantifying exposure to bright light of 250 mEDI lux and 1000 mEDI lux. Of note, we calculated differences in ten out of the 14 metrics analysed as differences in mean values of the metrics across the study. Thus, it could be that our results would differ when investigating differences in metrics for each day of the week or for specific times of the day. For example, in studies that examine the association between evening light exposure levels and health outcomes, filtering non-wear time using clusters of low illuminance might yield different metrics compared to calculating metrics on the raw data. In other words, longer recording periods might make the data more robust to artefacts and missing data, a finding which is supported by a recent preprint by Biller, Zauner and colleagues (2024), where multiple hours of missing daily data did not significantly influence average light exposure metrics for a dataset that spans a whole month. However, as more studies will need to confirm the generalisability of these findings in different, larger cohorts and in studies of varying duration, we recommend including non-wear detection as part of pre-processing pipelines and hereby present an easy-to-implement algorithm that can serve this purpose.

### Towards proposed best practices for collecting and reporting non-wear data

In the current study, participants had to perform three actions to report non-wear: press a button (event marker) on the device, place it in a black bag, and log this action in an app-based Wear log. We recognised that this could significantly burden participants and potentially result in lower data quality and implemented various countermeasures to minimise this risk. In this paragraph, we outline the strategies which we found to be successful.

Since the light glasses were very visible on participants’ faces, thus potentially contributing to more instances of non-wear, transparency about the device form factor was essential in exchanges with participants before screening. During the screening, emphasising the importance of wearing the device as much as possible and logging non-wear with honesty was successful in ensuring compliance. Our Wear log questionnaire provided options for retrospective logging of non-wear events as well as indicating where the device was located in case no black bag was used. Although these are scenarios impacting data quality, they are inevitable in ambulatory studies, and it is recommended to account for them and systematically label these instances rather than disregarding their occurrence (Van Der Donckt et al., 2024). This proactive and informed approach allows researchers to make better choices of how to handle this data in the analyses. Furthermore, providing participants with the rationale behind performing specific non-wear behaviour was important for compliance. Specifically, we explained that the use of the black bag would lead to low light levels, and that this would help our analyses. Maintaining contact with the participants during the week, monitoring their compliance, and reinstructing them when needed were essential to promote engagement throughout the study. This included continuous inspection of wear behaviour and interference when a Wear log entry appeared inconsistent with the previous entry. We acknowledge that this continuous monitoring can increase experimenter burden and does not scale well to studies with many participants. We recommend providing study participants with non-wear instruction sheets that can be directly consulted by participants in case of doubts on how to log a wear behaviour and even unexpected behaviour of the light logger. While a comprehensive review of challenges and solutions to promote accuracy of non-wear data collection in ambulatory light logging field studies is out of the scope of this paper, we refer the reader to related work in the field of actimetry, where relevant strategies to mitigate data entry challenges and ensure participant compliance have been examined (Balbim et al., 2021; Sriram et al., 2009; Van Der Donckt et al., 2024).

### Limitations

We acknowledge several limitations of the current study. First, we consider the self-reported information from the Wear log as ground truth. We acknowledge that self-reports may not be reliable, and they do not represent an optimal ground truth of when the light glasses were worn or not. While we tried our best to limit inconsistencies in the self-reported data by performing continuous data monitoring of Wear log entries and systematic pre-processing of this data, we ultimately cannot determine whether these entries represent participants’ actual non-wear behaviour. Second, our collection approach and our pre-processing pipeline have to be interpreted in light of our study sample, which consisted of healthy, compliant, and engaged participants. The high compliance of our participants also shows in the low number of Wear log entries that had to be cleaned (2.76% of all Wear log entries, due to for example open intervals, incorrect timestamps and duplicate entries). In large scale studies or clinical populations, continuously monitoring data and asking participants to report non-wear time as performed in the current study is not possible, and number of incorrect entries and open Wear log intervals would be higher than the one encountered in this study. Therefore, more extensive pre-processing pipelines might be required to ensure data quality and accurate detection of non-wear in larger studies and in studies with a low-compliance sample. As a consequence, we can consider the results of this study as the best possible outcome. Third, feasibility and accuracy of non-wear collection, detection, and handling strategies depend on specific characteristics of the light logger used in ambulatory studies. For example, for wrist-worn light loggers, skin temperature and conductance can be used as inputs to automated non-wear detection, and existing algorithms might be more useful than the approach taken here (Van Der Donckt et al., 2024). On the other hand, for chest and eye-level light loggers, different inputs are required for automated non-wear detection, and the strategies outlined in the current paper might be useful to further develop suitable algorithms for each light logger form factor. Fourth, our analysis on the comparison of light exposure metrics across the raw and non-wear corrected datasets is limited by our relatively small sample size, and in that differences are examined considering only one metric for each participant across the week, rather than looking at differences in daily light exposure metrics values. Lastly, the current study had a high temporal resolution with a measurement frequency of 10 seconds. Smoothing or re-sampling of light data might affect the performance of automated non-wear intervals detection, particularly for those of short duration. While more research is required on how to best pre-process light data and whether smoothing techniques affect non-wear detection, researchers should provide detailed information on how they apply smoothing transformations prior to non-wear detection on their datasets.

## Conclusion

The emergence of wearable light loggers in recent years has advanced the study of light exposure in free-living conditions. However, there remains a need of standardisation in pre-processing the large time series data produced by these devices, including the detection of non-wear intervals. In this study, we systematically collected non-wear data using multiple independent sources and continuously monitored participant’s self-reported non-wear behaviour. This approach provided a highly accurate non-wear ground truth, enabling a robust assessment of data-driven detection methods. Our findings demonstrate that, in a high compliance cohort, using a black bag for non-wear collection combined with an algorithm to detect resulting clusters of low illuminance is a promising strategy to manage non-wear time. Additionally, individual-level analysis of algorithm performance for low activity and low illuminance clusters suggests that combining activity and illuminance data could further improve non-wear detection. We believe that the methods and insights presented in the current study will inform the development of more advanced automated non-wear detection, including machine learning approaches. To support this effort, we are making our dataset available and hope to promote continued discussion to establish best practices for data pre-processing in ambulatory light exposure research.

## Declarations

### Ethics approval

This study was performed in line with the principles of the Declaration of Helsinki. Approval was granted by the Ethics Committee of the Technical University of Munich (TUM Ethics Committee 2023-115-S-KK).

### Consent to participate

Participants signed an informed consent prior to participation in this study.

### Consent for publication

Participants consented to publication of the data from this study by signing an informed consent.

### Availability of code, data and materials

All code and data analysed in this study are publicly available on GitHub (https://github.com/tscnlab/GuidolinEtAl_BehavResMethods_2025) under the MIT Licence (code) and CC-BY Licence (data).

### Authors’ contributions

Conceptualisation: CG, MS

Data curation: CG

Formal analysis: CG

Funding acquisition: MS

Methodology: CG, JZ, SLH

Project administration: CG

Software: CG, SLH

Resources: - Supervision: MS

Validation: - Visualisation: CG

Writing – original draft: CG

Writing – review & editing: CG, JZ, SLH, MS

### Open practice statement

Data and analysis code are available on GitHub at: https://github.com/tscnlab/GuidolinEtAl_BehavResMethods_2025. This study was not pre-registered.

### Declaration of AI-assisted technologies in the writing process

During the preparation of this work, the authors used ChatGPT to assist in reformulating sentences and improving the clarity of the manuscript. All content was subsequently reviewed and revised by the authors, who take full responsibility for the final version of the publication.

## Acknowledgments

We thank an anonymous reviewer for their suggestion to include equivalence testing as part of our analytic strategy.

