## Supplementary Materials for "Collecting, detecting and handling non-wear intervals in longitudinal light exposure data"

**Figure S1**

Complete data pre-processing pipeline adopted to clean and transform Wear log entries (ground truth)

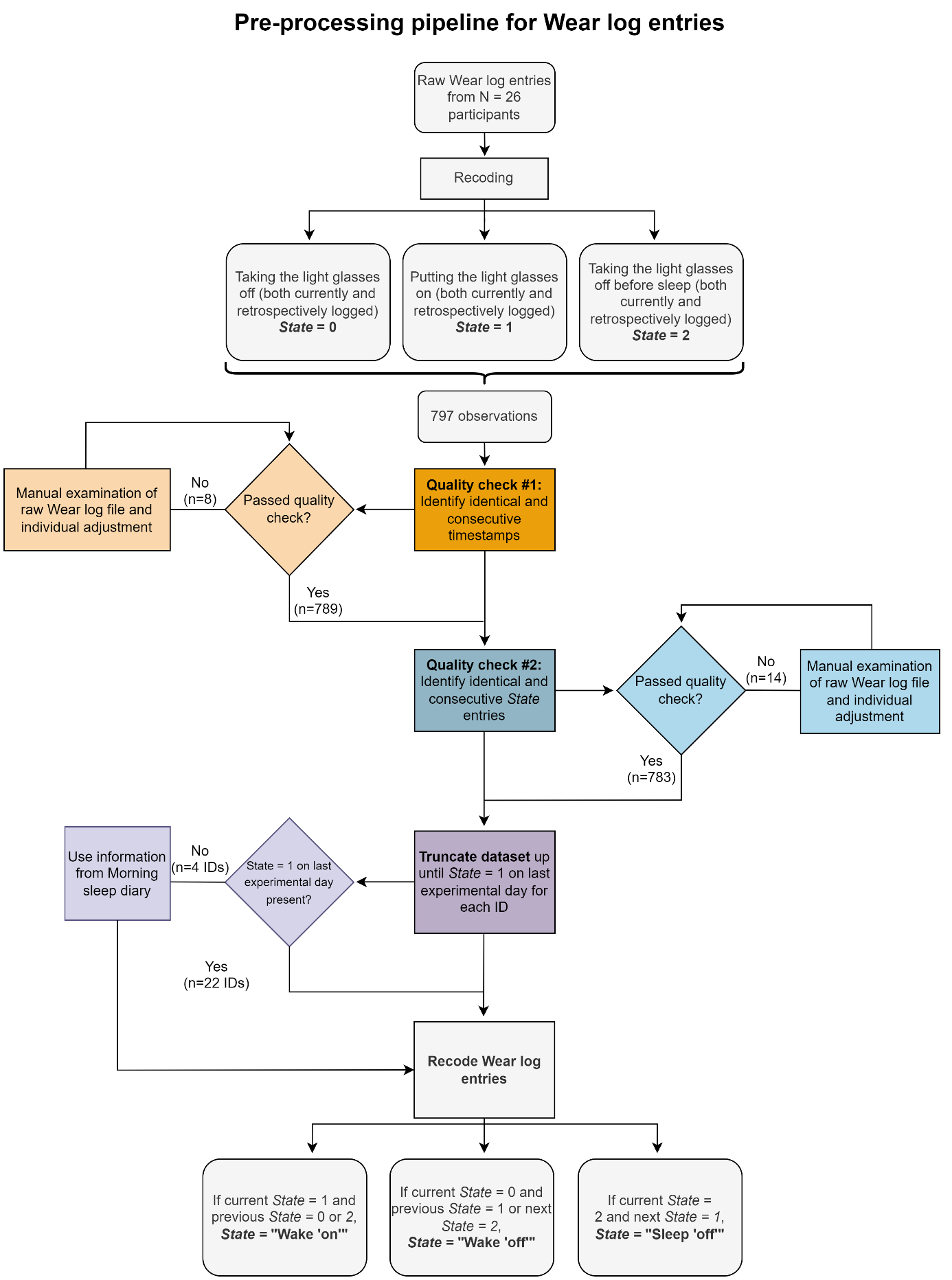

*Note.* Rectangles indicate processes, rhombuses are used for yes/no branching, and rectangles with smoothed corners are used for descriptions.

### Study materials: details on Firmoo.de glasses models

All non-prescription glasses were purchased with the following specifications: transparent glass type, standard (spherical) lens thickness, no coating. Furthermore, the following models were purchased:

- Size S: DBSN62240 model
- Size M: P181108R model
- Size L: TR31769 model

### Continuous data quality assurance: details on invalid entries

The following entries were considered invalid:

1. Consecutive Wear log entries of the same type
    *Example*: Two “Taking the light glasses off” entries were reported consecutively, without a “Putting the light glasses on” entry between them.
2. Inconsistent Wear log entries
    *Example*: Missing “Taking the light glasses off before sleep and placing them on a nightstand or flat surface” entry on a given day, but subsequent “Putting the light glasses on” entry.
3. Consecutive Wear log entries of different types but with the same timestamp *Example*: A “Taking the light glasses off” entry was logged at 13:33, and “Putting the light glasses on” entry was also logged at 13:33.

After identification of these entries, the participant was contacted via email and asked for clarification. Once the invalid entry was clarified with the participant, the entry was adjusted as follows:

- For cases 1 and 2 above, the participant was asked to log the missing entry as a “retrospective” event on the Wear log.
- For case 3, the researcher noted this down in the participant’s spreadsheet.

### Data cleaning of Wear log entries: details on manually adjusted entries

#### Details for Quality check #1

The eight entries not passing Quality check #1 were adjusted as follows:

1. If two consecutive entries had identical *Datetime*, but different *State*, the automated timestamp of the *startdate* column in the raw data was consulted. This is an automated timestamp created by the MyCap app when participants log any event in the questionnaire. If the timestamp of the *startdate* was different to that of *Datetime*, the *startdate* timestamp was considered the correct, valid timestamp for the entry in question. The value of *Datetime* was modified manually to this value (n=4).
2. If two consecutive entries had identical *Datetime* and *State*, and the timestamp of the *startdate* column in the raw data coincided with *Datetime*, only the first Wear log entry was kept, and the second was considered accidental and thus removed manually (n=2).
3. If two consecutive entries had identical *Datetime*, but different *State*, and the timestamp from the *startdate* column coincided with *Datetime*, careful manual inspection of the raw wear log file was performed. This revealed the two following instances:
1. The participant had two legitimate *State* changes within the same minute, meaning that they spent less than one minute in a single *State,* leading to consecutive identical *Datetime* for two different consecutive *State* values. To correct for this, 60 seconds were added manually to the second *Datetime* (n=1).
2. The participant initially logged a ‘Taking the light glasses off’ close to their bedtime (as indicated in the sleep diary), and then logged a ’Taking the light glasses off before sleep and placing them on a nightstand or a flat surface’ immediately after the initial log. Only the latter log was kept (n=1).

#### Details for Quality check #2

The 14 entries not passing Quality check #2 were manually adjusted as follows:

1. If two consecutive entries had the same *State* and one was ‘retrospective’ and the other one ‘current’, only the current entry was kept, under the assumption that ‘current’ entries are more accurate than ‘retrospectively logged’ entries (n=2).
2. If three consecutive entries had the same *State*, visual inspection of participants’ light and activity data within this time window was performed. If either light or activity data suggested that these three *State* were not the same, then the "middle" entry was considered accidental entry and changed accordingly (n=1).
   - Detailed description: Three consecutive *State* = 0 (“Taking the light glasses off”) entries at 19:45, 19:54 and 20:14, but light and activity data suggested, by visual inspection, non-wear between 19:45 and 19:54, and a wear period between 19:54 and 20:14. 19:54 was manually adjusted to *State* = 1.
3. If participants logged a “Taking the light glasses off before sleep and placing them on a nightstand or flat surface” entry after midnight but forgot to update the date to the "new" day, the date was manually corrected to the “new” date (n=3).
4. If two different entries on the app were logged as the same entry in the wear log, this was detected as technical problem and visual inspection of light values within the two time entries helped assign each entry to a correct timestamp (n=1).
5. If a "Putting the light glasses” entry in the morning was missing entirely, the timestamp of *Datetime* was assigned according to information from the “Morning sleep diary” (*out_ofbed* variable). This questionnaire contains timestamps for the events “waking up” and “going to sleep” and was thus assumed to be the closest approximation for this missing wear log entry (n=1).
6. If an erroneous entry occurred on the last experimental day (Monday) after the first *State* = 1, it was deleted, since not relevant to the analysis (n=1).
7. If an entry failing Quality check #2 was linked to another erroneous entry failing the same check, and adjusting the latter automatically resolved the former, this was left unadjusted (n=3).

**Figure S2**

Precision recall curves for each input parameter of the cluster detection algorithm for low illuminance

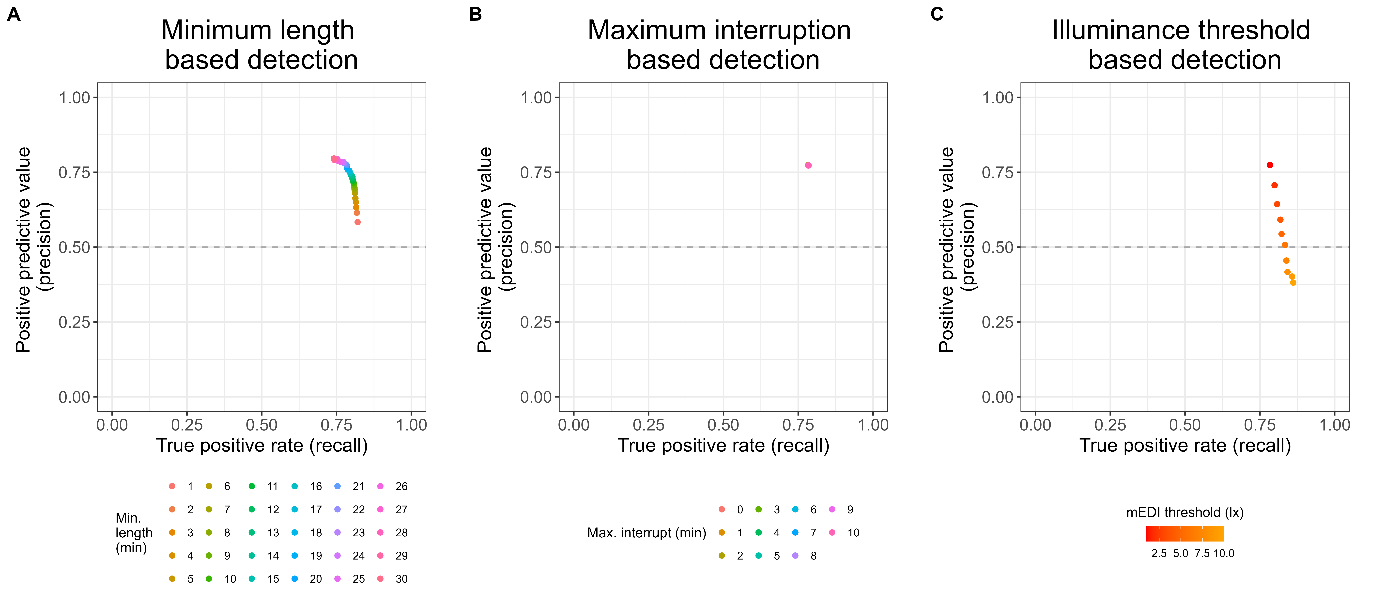

*Note.* (A) Precision recall curve for detection of low illuminance clusters with varying clusters minimum lengths, with mEDI threshold <1 lux and maximum interruption length=0 minutes. (B) Precision recall curve for detection of low illuminance clusters with varying maximum interruption lengths, with mEDI threshold <1 lux and minimum cluster length=21 minutes. (C) Precision recall curve for detection of low illuminance clusters with varying mEDI thresholds (1 to 10 lux at 1-step increments), with minimum cluster length=21 minutes and maximum interruption length=0 minutes.

**Figure S3**

Precision recall curves for each input parameter of the cluster detection algorithm for low activity

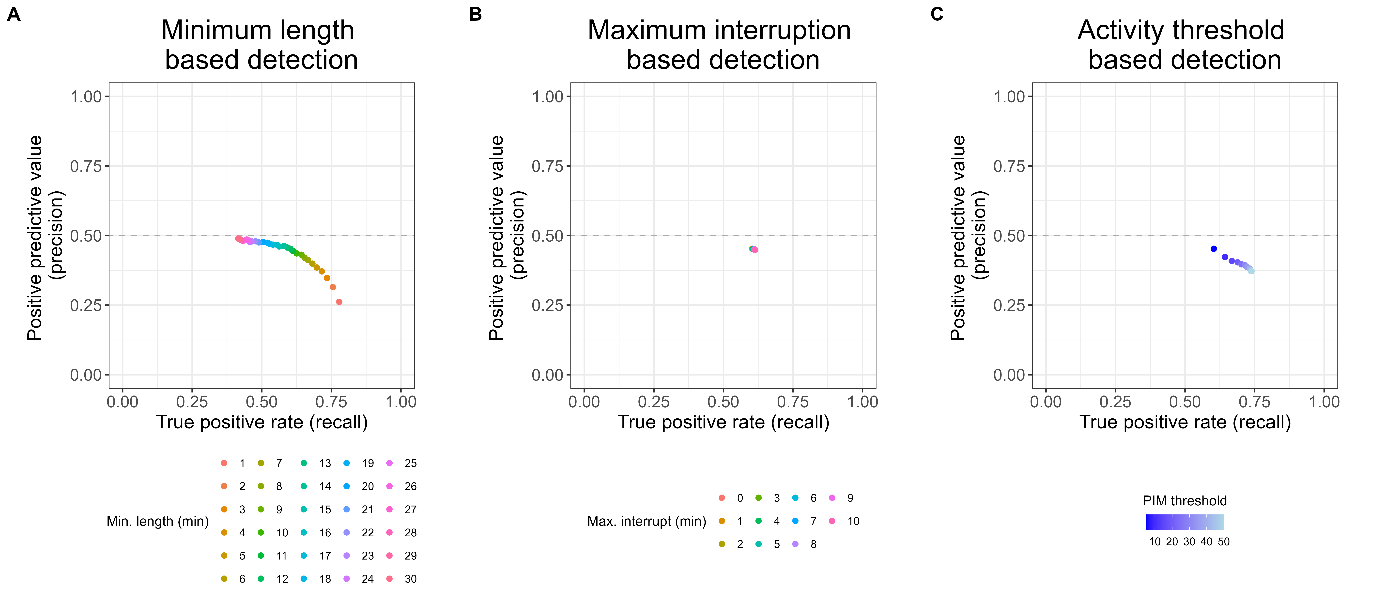

*Note.* (A) Precision recall curve for detection of low activity clusters with varying clusters minimum lengths, with PIM threshold <5 and maximum interruption length=0 minutes. (B) Precision recall curve for detection of low activity clusters with varying maximum interruption lengths, with PIM threshold <5 and minimum cluster length=12 minutes. (C) Precision recall curve for detection of low activity clusters with varying PIM thresholds (5 to 50 lux at 5-step increments), with minimum cluster length=12 minutes and maximum interruption length=0 minutes.

**Figure S4**

F1 score as a function of minimum cluster length for activity and illuminance clusters

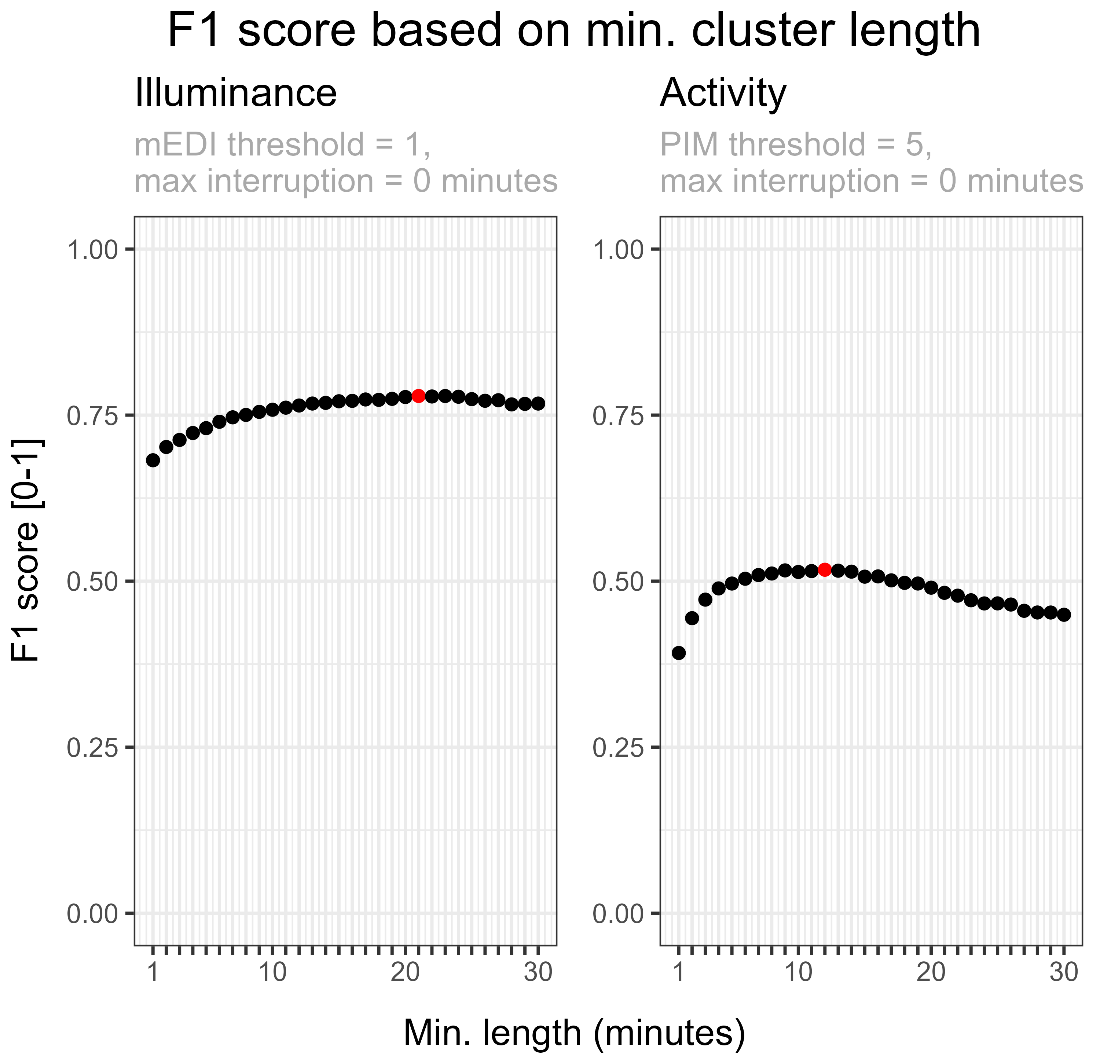

*Note.* F1 score plotted as a function of minimum cluster length considered for illuminance (left panel) and activity (right panel). Red dots indicate the highest F1 score: for illuminance, this is 0.78, corresponding to a minimum length of 21 minutes, for activity, this is 0.52, corresponding to a minimum length of 12 minutes.

**Figure S5**

Difference between self-reported “Sleep ‘off’” time (Wear log) and sleep time (sleep diary) entries.

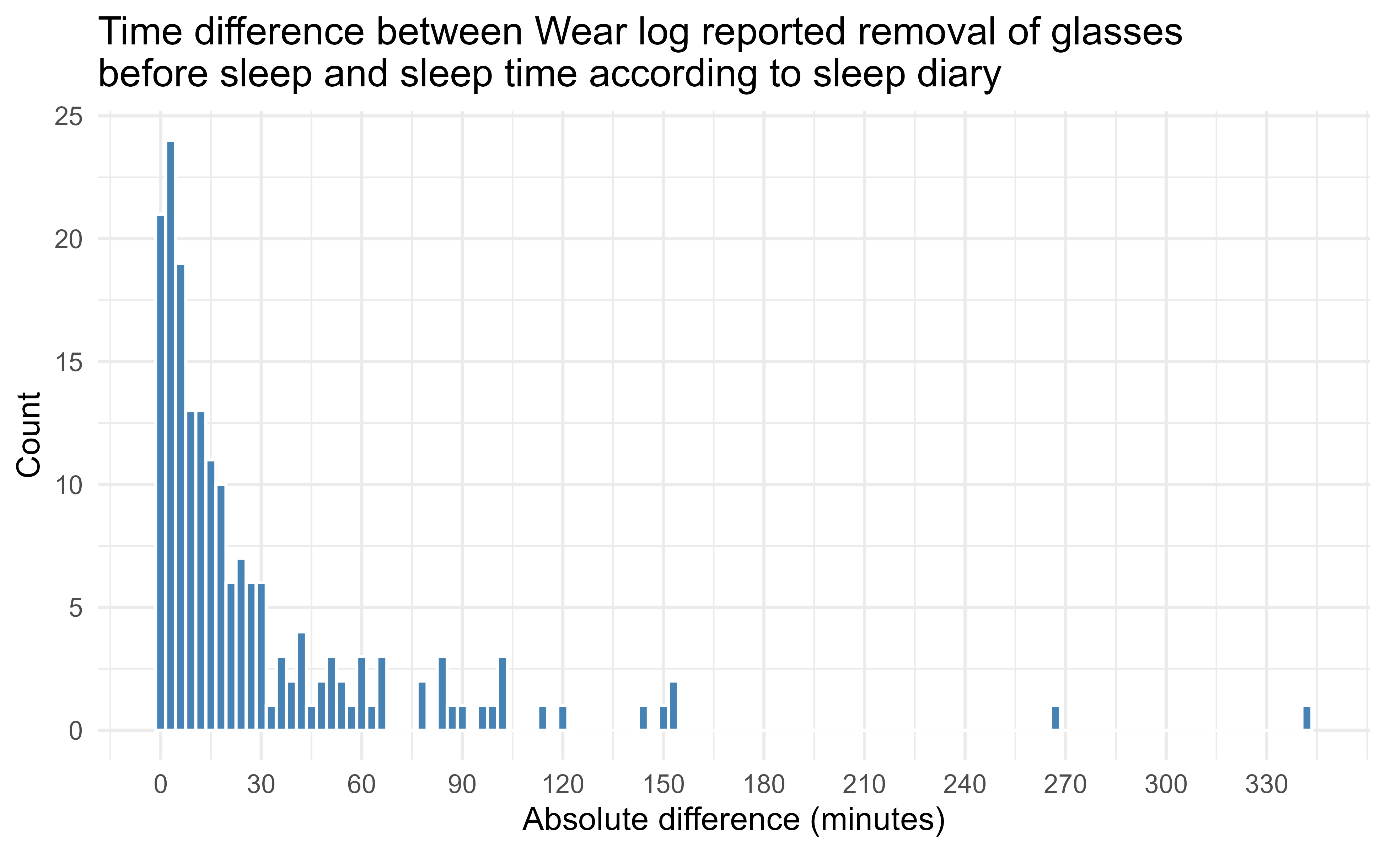

*Note*. Sleep time was defined as the time when the participants tried to fall asleep after going to bed.

**Figure S6**

Precision recall curves for low illuminance clusters based on daytime and nighttime.

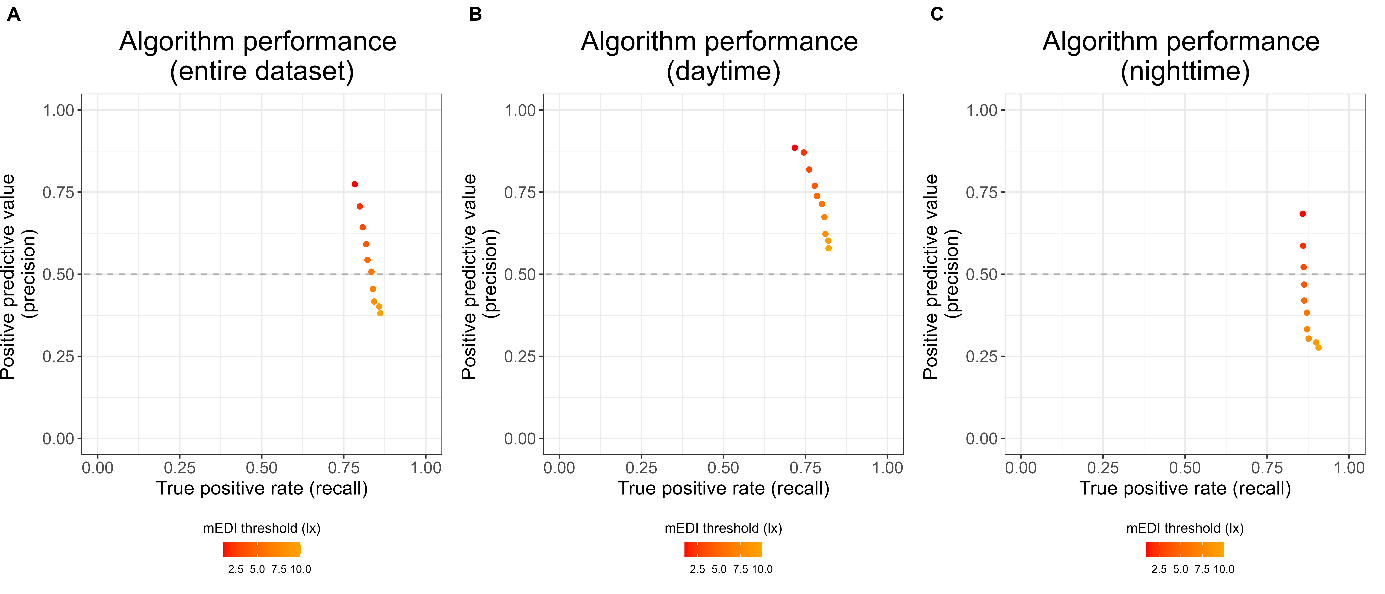

*Note.* A minimum cluster length of 21 minutes and a maximum interruption length of 0 minutes were used as input parameters. The dataset was split into daytime and nighttime based on the photoperiod information from the coordinates where the study was conducted (Tübingen, Germany: 48.5216° N, 9.0576° E). (A) Algorithm performance for the entire dataset; (B) algorithm performance for daytime periods only; (C) algorithm performance for nighttime periods only.

**Figure S7**

Precision recall curves for low activity clusters depending on the activity-quantifying parameter (PIM, TAT and ZCM)

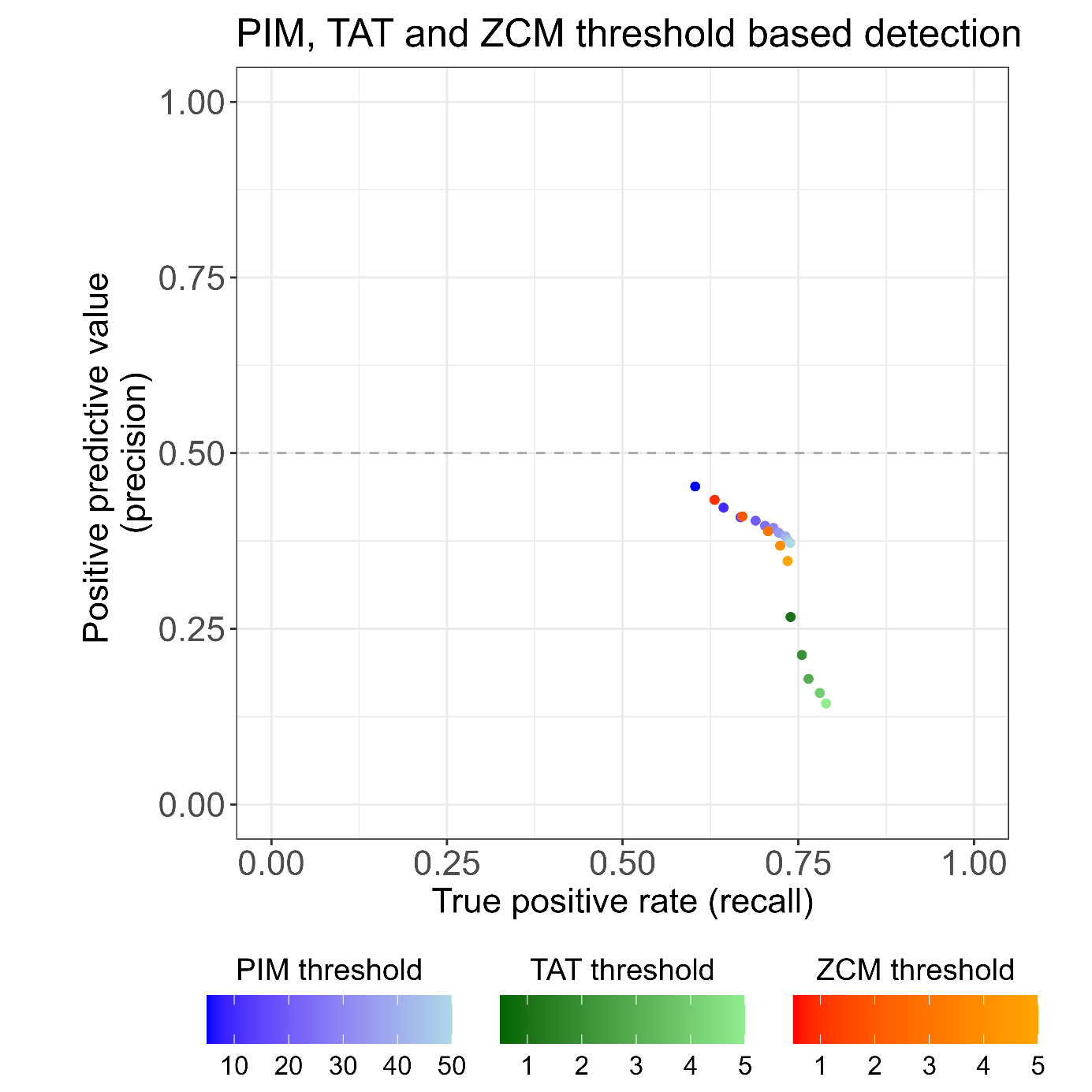

*Note.* A minimum cluster length of 12 minutes and a maximum interruption length of 0 minutes were used as input parameters for all three algorithms. Activity thresholds were from 5 to 50 (at 5-step increments) for PIM, and from 0.5 to 5.0 (at 0.5-step increments) for TAT and ZCM, to reflect small values of each parameter.

**Figure S8**

Cluster detection algorithm performance for low activity depending on pre-processing of activity values

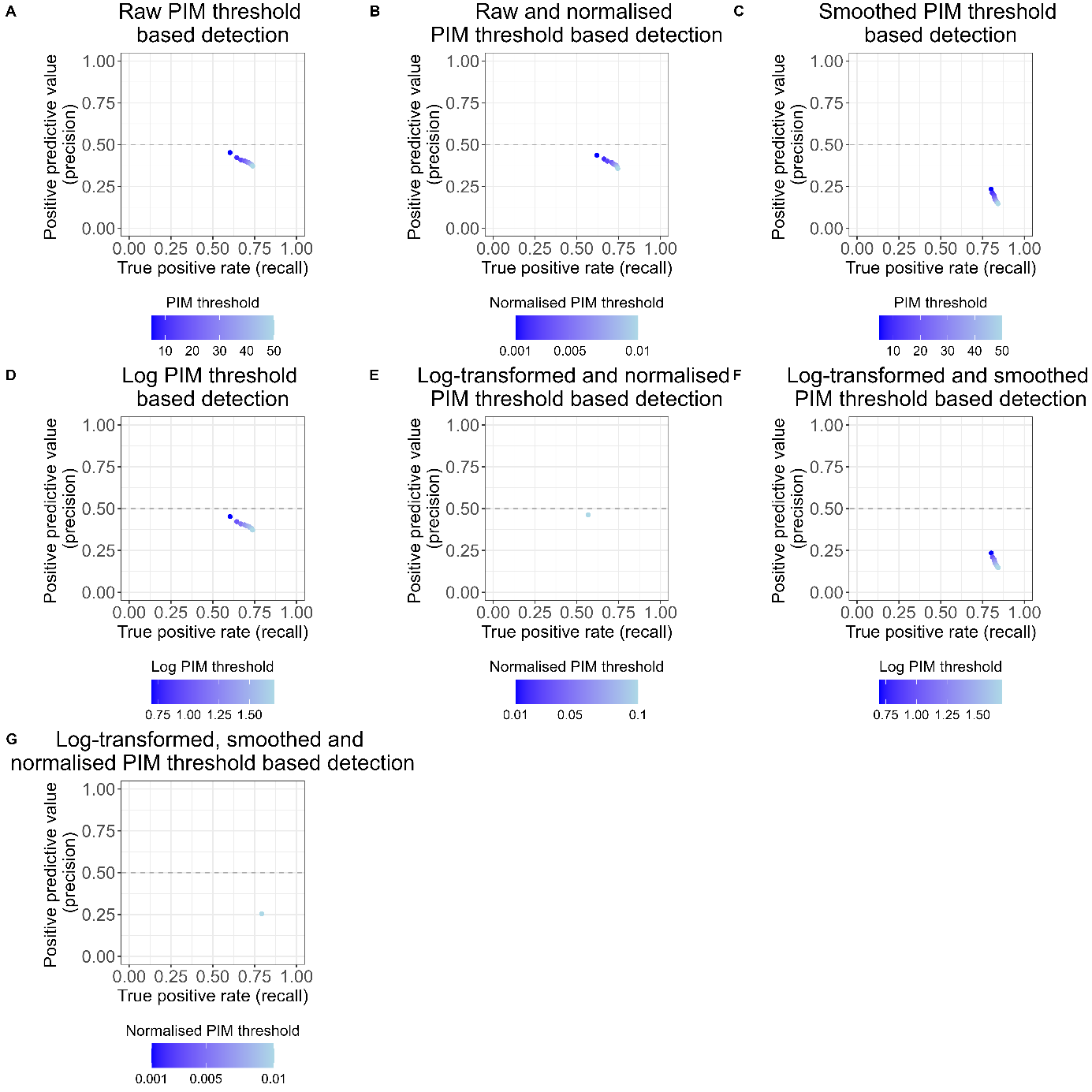

*Note.* (A) Precision recall curve for detection of low activity clusters from raw PIM values (F1=0.52). (B) Precision recall curve for detection of low activity clusters from normalised PIM values, using min-max normalisation (F1=0.51). (C) Precision recall curve for detection of low activity clusters from smoothed PIM values (F1=0.36). Smoothing was performed using a rolling median with window of 10 minutes. (D) Precision recall curve for detection of low activity clusters from log10 transformed PIM values (F1 = 0.52). (E) Precision recall curve for detection of low activity clusters from log10-transformed and normalised (min-max normalisation) PIM values (F1=0.51). Note that regardless of the threshold, the F1 score is the same. (F) Precision recall curve for detection of low activity clusters from log10-transformed PIM and then smoothed PIM values (F1=0.36). Smoothing was performed using a rolling median with window of 10 minutes. (G) Precision recall curve for detection of low activity from log10 transformed, smoothed and normalised PIM values (F1=0.38). Smoothing was performed using a rolling median with window of 10 minutes, and min-max normalisation was used. Note that regardless of the threshold, the F1 score is the same. A minimum cluster length of 12 minutes and a maximum interruption length of 0 minutes were used as input parameters for all algorithms applied to the different pre-processed PIM values shown in this figure. In case of log-transformation, a value of 0.1 was added to the data prior to transformation in order to handle 0 values.

**Figure S9**

Bootstrap distributions for equivalence testing of post-hoc pairwise comparisons

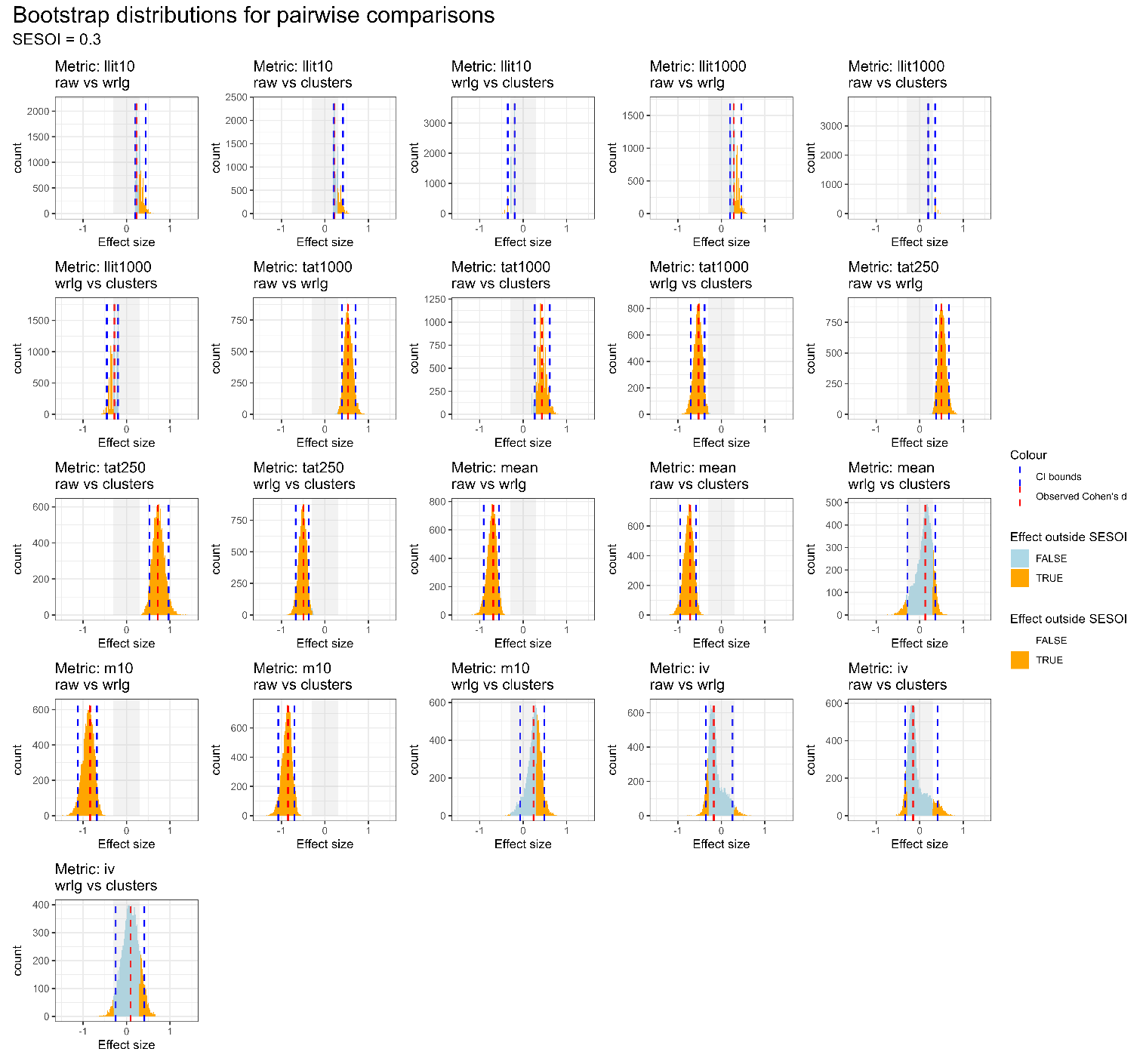

*Note.* Bootstrap distributions for each post-hoc pairwise comparison from a significant Friedman’s test. Each panel represents a pairwise comparison for a given metric. The smallest effect size of interest (SESOI) for the differences between two datasets was set to *d* = 0.3 (indicated by the grey shaded area). Confidence intervals and the observed Cohen’s d are represented by vertical blue and red dashed lines, respectively. Orange and light blue bars on the histogram represent bootstraps outside and inside the SESOI, respectively.

**Figure S10**

Proportion of bootstraps outside of the SESOI bounds based on SESOI benchmark decision.

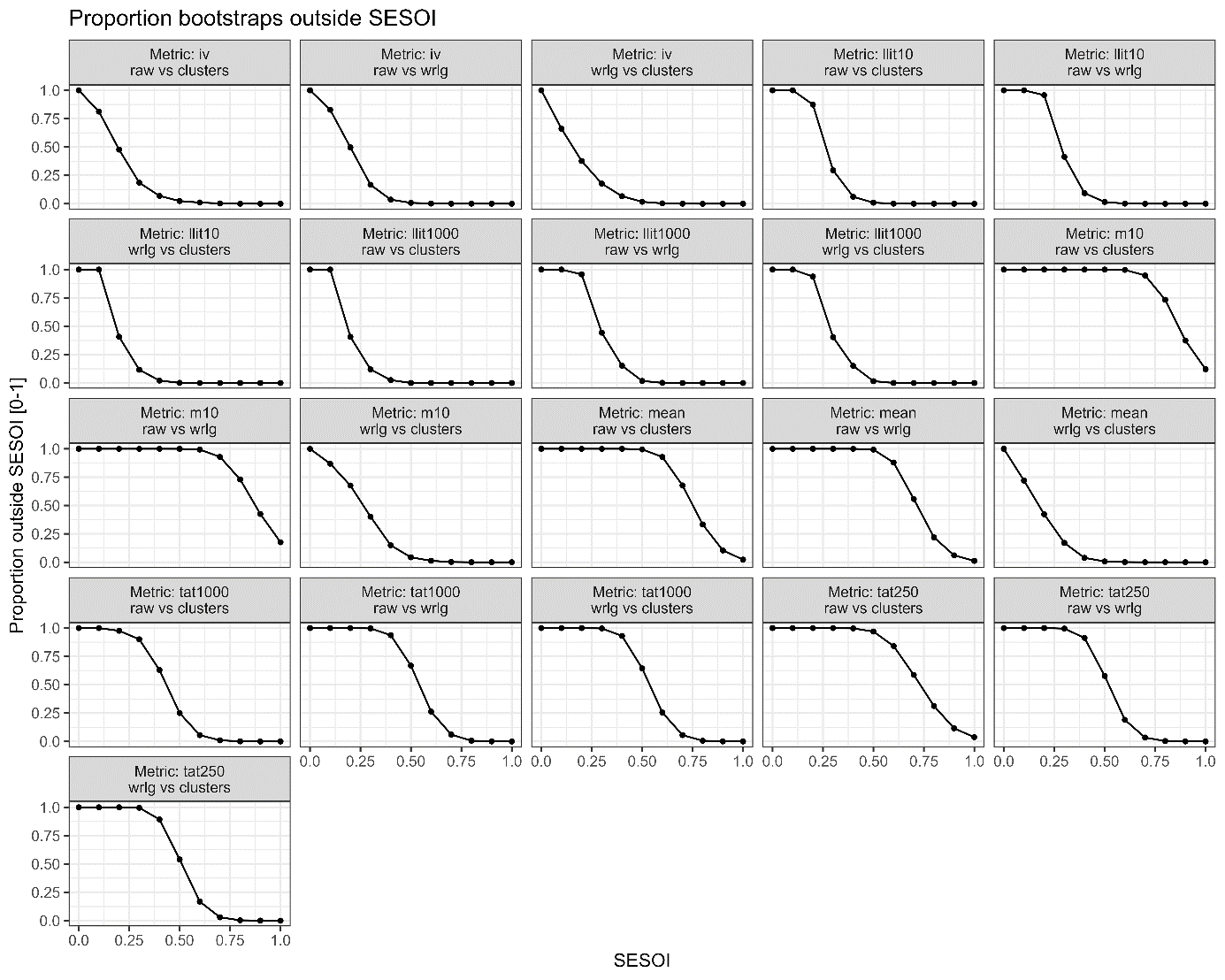

*Note.* The proportion of bootstraps outside of the SESOI bounds is shown for each post-hoc pairwise comparison as a function of SESOI decision. The X axis represents possible SESOI benchmarks (corresponding to *d* = 0.0 to *d* = 1.0 at 0.1 steps).

**Figure S11**

Equivalence test results based on SESOI benchmark decision

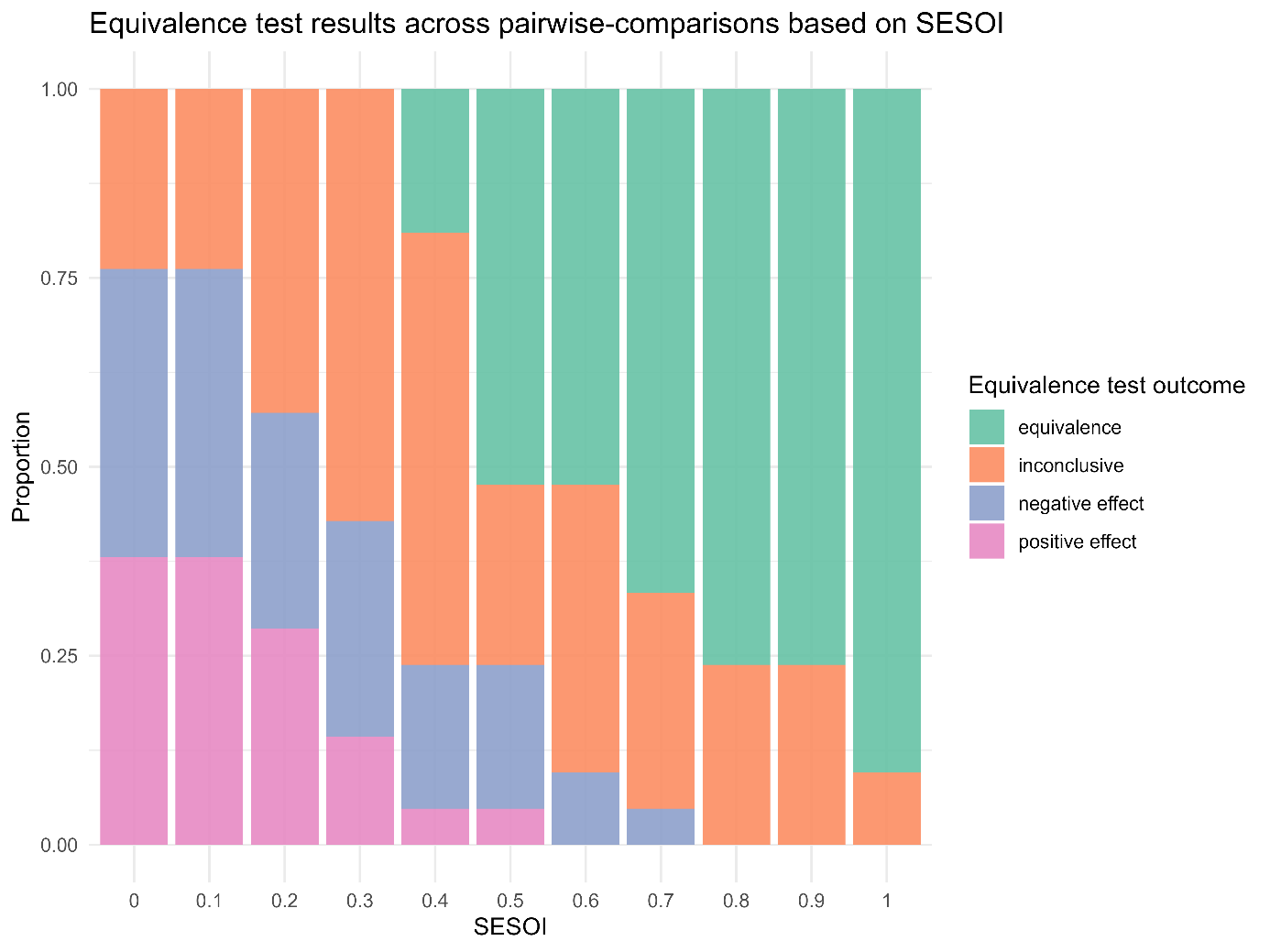

*Note.* Proportions for each possible outcome of equivalence testing are shown based on SESOI benchmark decision. Equivalence test outcomes include: equivalence (confidence intervals inside the SESI bounds), inconclusive (one or both confidence intervals extending beyond the SESOI bounds), meaningfully negative effect (both confidence intervals entirely below the lowest SESOI bound), meaningfully positive effect (both confidence intervals entirely above the upper SESOI bound).

**Table S1**

Friedman’s test results for n=14 light exposure metrics across three datasets

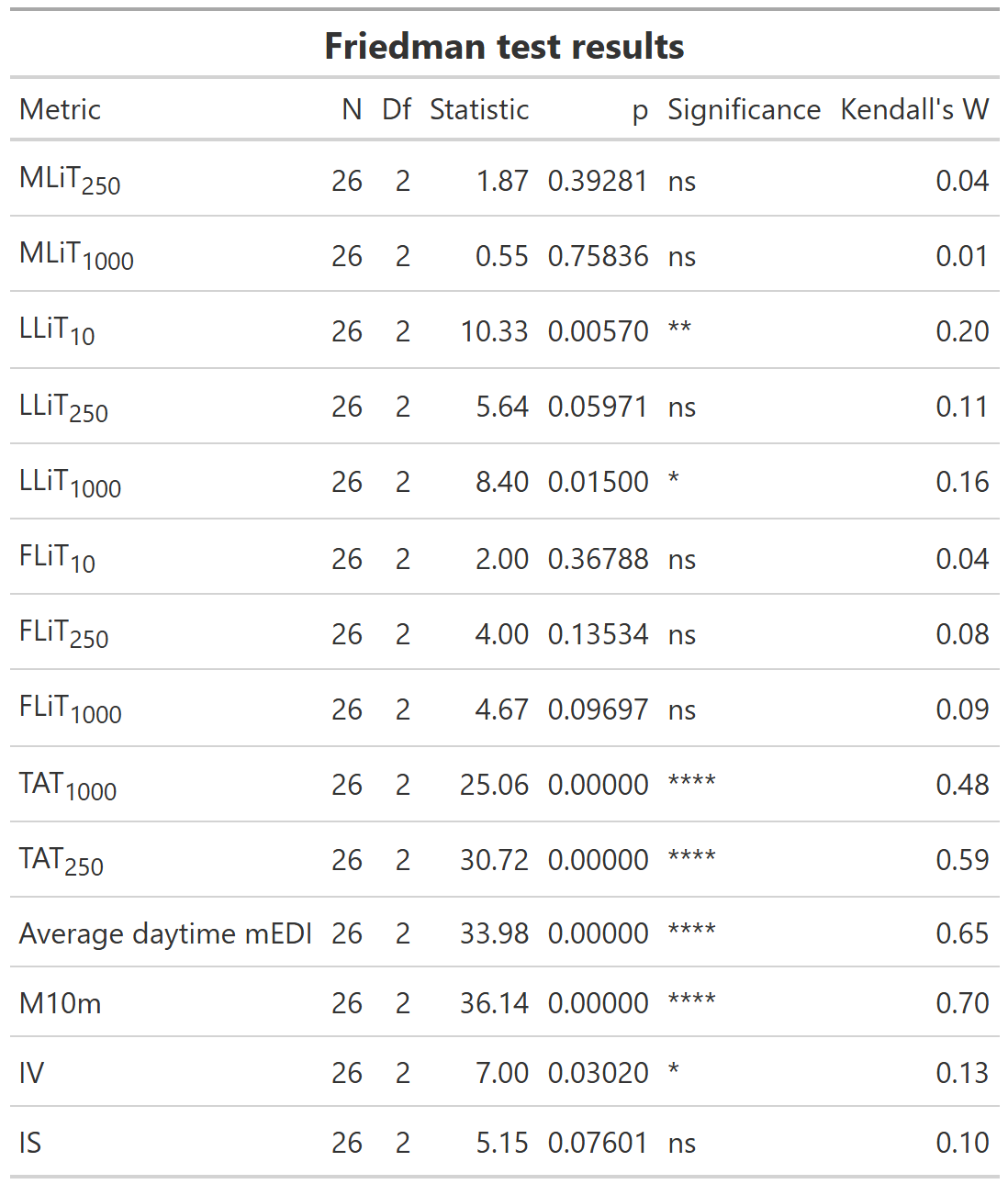

*Note.* Kendall’s W indicates the effect size of the difference (0-1). Significant values are shown at threshold p<0.05.

**Table S2**

Means and SDs of n=14 light exposure metrics

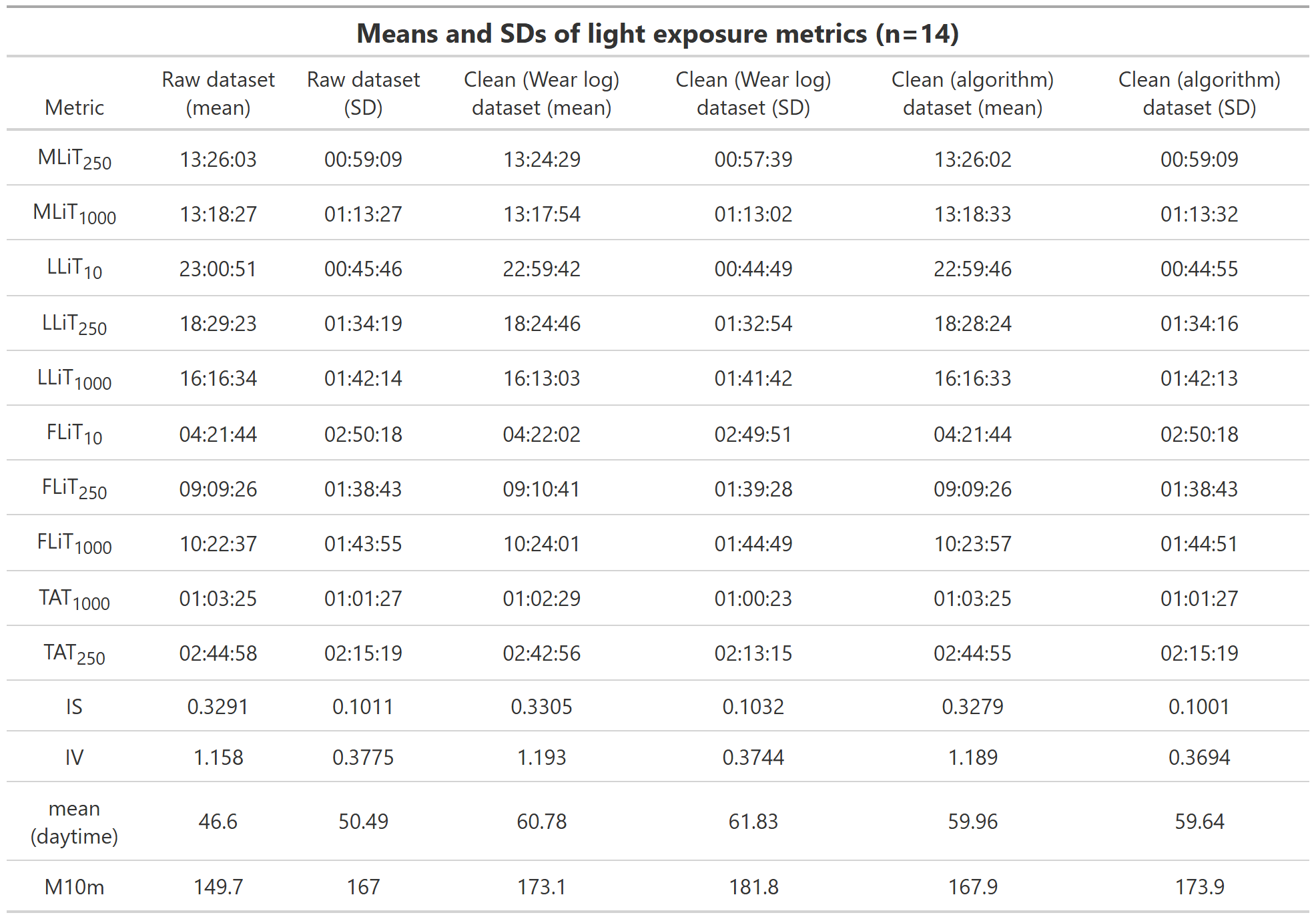
